# Overcoming myeloid-driven resistance to CAR T therapy by targeting SPP1

**DOI:** 10.1101/2025.04.01.646202

**Authors:** Sharareh Gholamin, Heini M. Natri, Yuqi Zhao, Shengchao Xu, Maryam Aftabizadeh, Begonya Comin-Anduix, Supraja Saravanakumar, Christian Masia, Robyn A. Wong, Lance Peter, Mei-i Chung, Evan D. Mee, Brenda Aguilar, Renate Starr, Davis Y. Torrejon, Darya Alizadeh, Xiwei Wu, Anusha Kalbasi, Antoni Ribas, Stephen Forman, Behnam Badie, Nicholas E. Banovich, Christine E. Brown

## Abstract

Chimeric antigen receptor (CAR) T cell therapy faces notable limitations in treatment of solid tumors. The suppressive tumor microenvironment (TME), characterized by complex interactions among immune and stromal cells, is gaining recognition in conferring resistance to CAR T cell therapy. Despite the abundance and diversity of macrophages in the TME, their intricate involvement in modulating responses to CAR T cell therapies remains poorly understood. Here, we conducted single-cell RNA sequencing (scRNA-seq) on tumors from 41 glioma patients undergoing IL13Rα2-targeted CAR T cell therapy, identifying elevated suppressive *SPP1* signatures predominantly in macrophages from patients who were resistant to treatment. Further integrative scRNA-seq analysis of high-grade gliomas as well as an interferon-signaling deficient syngeneic mouse model—both resistant to CAR T therapy—demonstrated the role of congruent suppressive pathways in mediating resistance to CAR T cells and a dominant role for *SPP1*+ macrophages. SPP1 blockade with an anti-SPP1 antibody abrogates the suppressive TME effects and substantially prolongs survival in IFN signaling-deficient and glioma syngeneic mouse models resistant to CAR T cell therapy. These findings illuminate the role of *SPP1*+ macrophages in fueling a suppressive TME and driving solid tumor resistance to CAR cell therapies. Targeting *SPP1* may serve as a universal strategy to reprogram immune dynamics in solid tumors mitigating resistance to CAR T cell therapies.

## Main

Chimeric antigen receptor (CAR) T cell therapy has yielded transformative success in hematologic malignancies, but its application in solid tumors has been met with mixed success^1^. Solid tumors exist within a highly structured yet dynamic microenvironment that dictates their growth, progression, and response to therapy. The tumor microenvironment (TME) can limit immune cell infiltration and foster treatment resistance through diverse, patient-specific mechanisms^2–4^. While some tumors exhibit immune activity that can support the efficacy of immunotherapy, others remain refractory, suggesting that response heterogeneity is influenced by distinct cellular programs within the TME^2,5^. However, the precise interactions within the TME that dictate CAR T cell response remain poorly understood, warranting further investigation into the regulatory networks that shape therapeutic outcomes in solid tumors.

These challenges are particularly pronounced in glioblastoma and other high-grade gliomas (GBM/HGGs), which are notorious for their immunosuppressive milieu and pronounced patient-specific heterogeneity^6^. In our recently reported phase 1 trial evaluating IL13Rα2-targeted CAR T cells in recurrent GBM/HGGs, we observed heterogeneous therapeutic responses. Improved outcomes were linked to higher infiltration of CD3+ T cells in pre-treatment tumors and increased levels of IFN-pathway cytokines during therapy^7^. While disease control was achieved in half of the patients (stable disease or better), the other half progressed through treatment, thereby highlighting ongoing challenges that remain for addressing the resistance of GBM/HGG. This resistance, in part, is driven by the inherent heterogeneity in intratumoral immune networks and the prevalence of suppressive programs governed by abundant heterogeneous macrophages.

The distinct biology of GBM/HGGs is shaped by a confluence of tumor-intrinsic properties and the profoundly suppressive TME. This complex ecosystem is defined by spatial and cellular heterogeneity, where immune, stromal, and metabolic interactions converge to drive tumor progression and therapeutic resistance^8–10^. While the abundance of myeloid components in GBM/HGGs is well-recognized, their definitive role in mediating resistance to cellular therapy remains poorly understood^11^. Moreover, the multipolarity and plasticity of macrophages within individual tumors and across patients add further complexity to their contribution to immune suppression and therapeutic failure. These unresolved questions highlight the urgent need for deeper investigation into the immune landscape of GBM/HGGs to inform strategies that can overcome the barriers imposed by the TME and improve responses to CAR T cell therapy.

### Tumor microenvironment features shape clinical response to CAR T cell therapy in GBM/HGGs

Recognizing the unique challenges posed by solid tumors to CAR T cell therapy, we set out to identify dominant networks within GBM/HGG associated with patient outcomes to adoptive T cell therapy. To delineate characteristics of the GBM/HGG microenvironment influencing responsiveness to CAR T cell therapy, we performed scRNA-seq on freshly dissociated pretreatment tumors from our recent clinical trial evaluating IL13Rα2-targeted CAR T cell therapy (Fig. 1a, Extended Fig. 1a, Supplementary Table 1)^7^. Out of 58 response-evaluable patients, 41 had sufficient tumor samples for scRNA-seq analysis. Patients were stratified based on treatment outcome, comparing patients who progressed through therapy (progressive disease abbreviated hereon PD; n=23) to those who had a more favorable response of stable disease or better (stable disease, SD, partial response, PR, and complete response, CR, abbreviated hereon as SD/CR; n=18) (Fig. 1b). Additionally, stratification was performed based on baseline CD3+ T cell infiltration, comparing tumors with low CD3+ T cell infiltration (CD3 IHC score 0, 1, and 2; CD3-low) to those with medium to high infiltration (CD3 IHC score 3 and 4; CD3-med/high). We have previously reported that GBM/HGG tumors with higher CD3+ T cell infiltration pre-CAR T treatment correlated with improved clinical outcomes (Fig. 1b)^7^. This dual stratification aimed to uncover shared pathways that could reveal universal features of the TME associated with GBM/HGG response to CAR T cell therapy. A semi-supervised lineage-based approach classified 51,829 cells as tumor, immune and stromal cell populations (Fig. 1c, Extended Fig. 1b,c). The 27,674 immune and stromal cells (2,072 fibroblasts, 10,518 lymphocytes, 15,084 myeloid cells) were further sub-clustered into 24 distinct populations (Fig. 1d, Extended Fig. 1d-f, Supplementary Tables 2-5) to interrogate microenvironment features.

**Figure 1.**
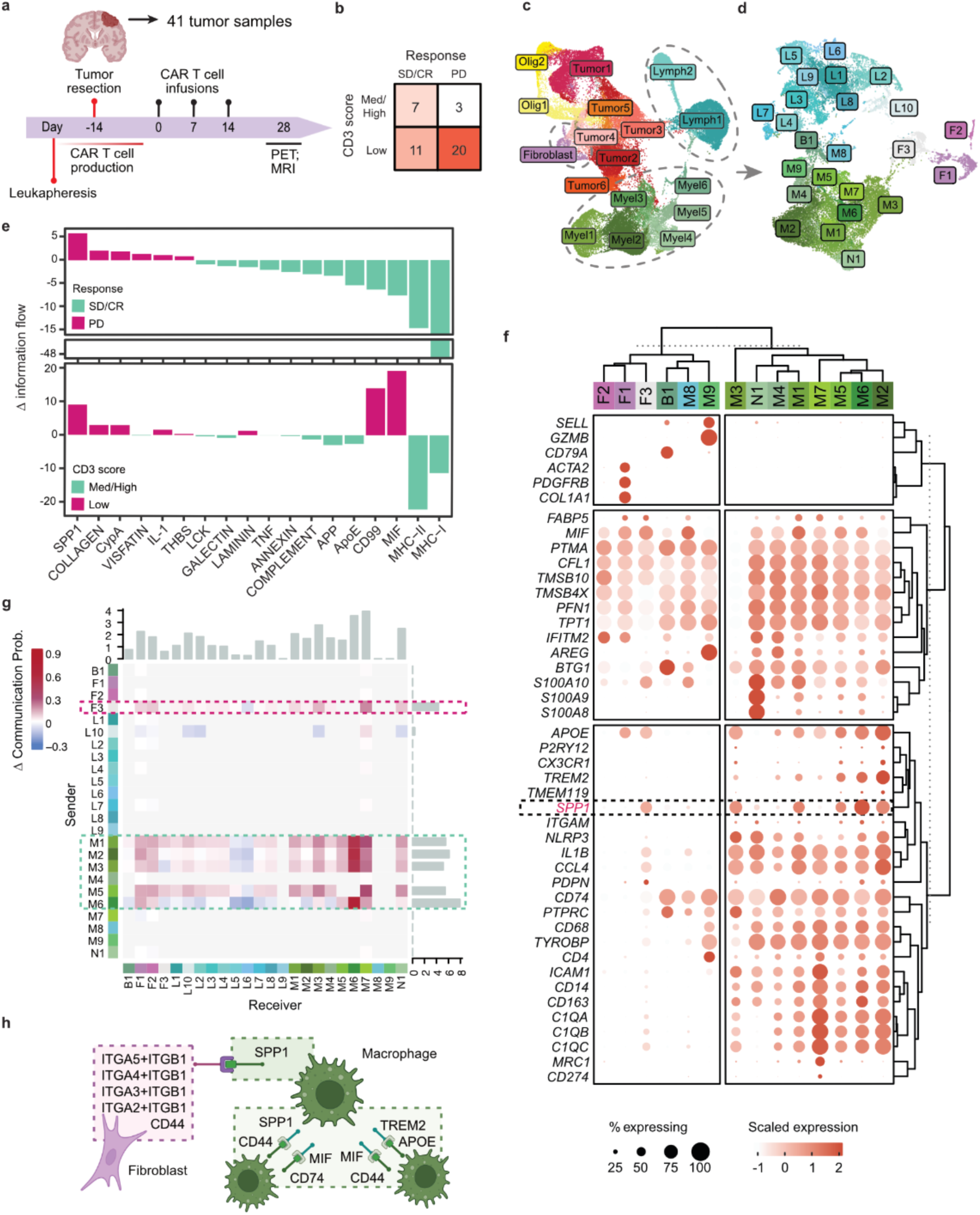
Gene expression profiling of GBM/HGGs reveals signaling pathways in the TME associated with response and resistance to IL13Rα2 CAR T therapy. **a**, A schematic illustration of the IL13Rα2 CAR T cell trial (NCT02208362) from which GBM/HGG tumor resection samples (n = 41) were collected for scRNA-seq analysis. **b**, Numbers of GBM/HGG tumor samples with medium/high (3-4) or low (0-2) CD3 score by IHC and samples that did (SD, PR and CR abbreviated going forward as SD/CR) or did not (PD) respond to IL13Rα2 CAR T therapy. **c**, UMAP dimensionality reduction of 51,829 cells from the whole tumor across 41 donors, pseudocolored by cluster identity. **d**, UMAP of 27,674 immune and stromal cells pseudocolored by cluster identity. **e**, Differentially regulated ligand-receptor signaling pathways in immune subclusters of the TME, comparing SD/CR vs. PD tumors (top) and CD3-med/high vs, CD3-low tumors (bottom). Significant pathways were selected with a two-sided Wilcoxon p < 0.01 in the comparison of SD/CR vs. PD, and pathways with an absolute delta of information flow > 0.8 were plotted, with positive values indicating increased signaling in PD or CD3-low tumors. The information flow for a given signaling pathway is defined by the sum of communication probability among all pairs of cell groups in the inferred network. **f**, Expression of marker features among the immune and stromal cells and expression status of discriminating features for each cluster depicted in (d). **g**, Differential *SPP1* signaling interaction strength between all pairs of analyzed cell types in a comparison of SD/CR and PD tumors. Positive values indicate increased signaling in PD tumors. **h**, Schematic of the top three cell-cell communication pathways in PD tumors compared to SD/CR tumors.

To identify pretreatment cellular communication networks underlying clinical response to CAR T therapy, we evaluated receptor-ligand interaction probabilities among known cell-cell communication networks by calculating the sum of communication probabilities among all pairs of cell types in the TME subsets as well as the whole tumor. In favorably responding patients, those who achieved SD/CR or had increased intra-tumoral CD3 infiltrates, the pretreatment TME signaling patterns were significantly different from non-responders (Fig. 1e, Supplementary Table 6). Specifically, MHC-presentation pathways, including both MHCI and MHCII, were up-regulated in responding patients (SD/CR and CD3-med/high; p < 0.01) when considering the whole tumor (Fig. 1c, Extended Fig. 2a) as well as the TME subset (Fig. 1d-e). Consistent with this observation, total lymphocyte proportions (Lymph1 and Lymph2) were significantly increased in SD/CR compared to PD (Extended Fig. 2b,c). These results align with improved patient outcomes following CAR T cell therapy being associated with higher pretreatment intratumoral T cell levels, as assessed by immunohistochemistry, and increased IFN-signaling, as previously reported^7^.

**Figure 2.**
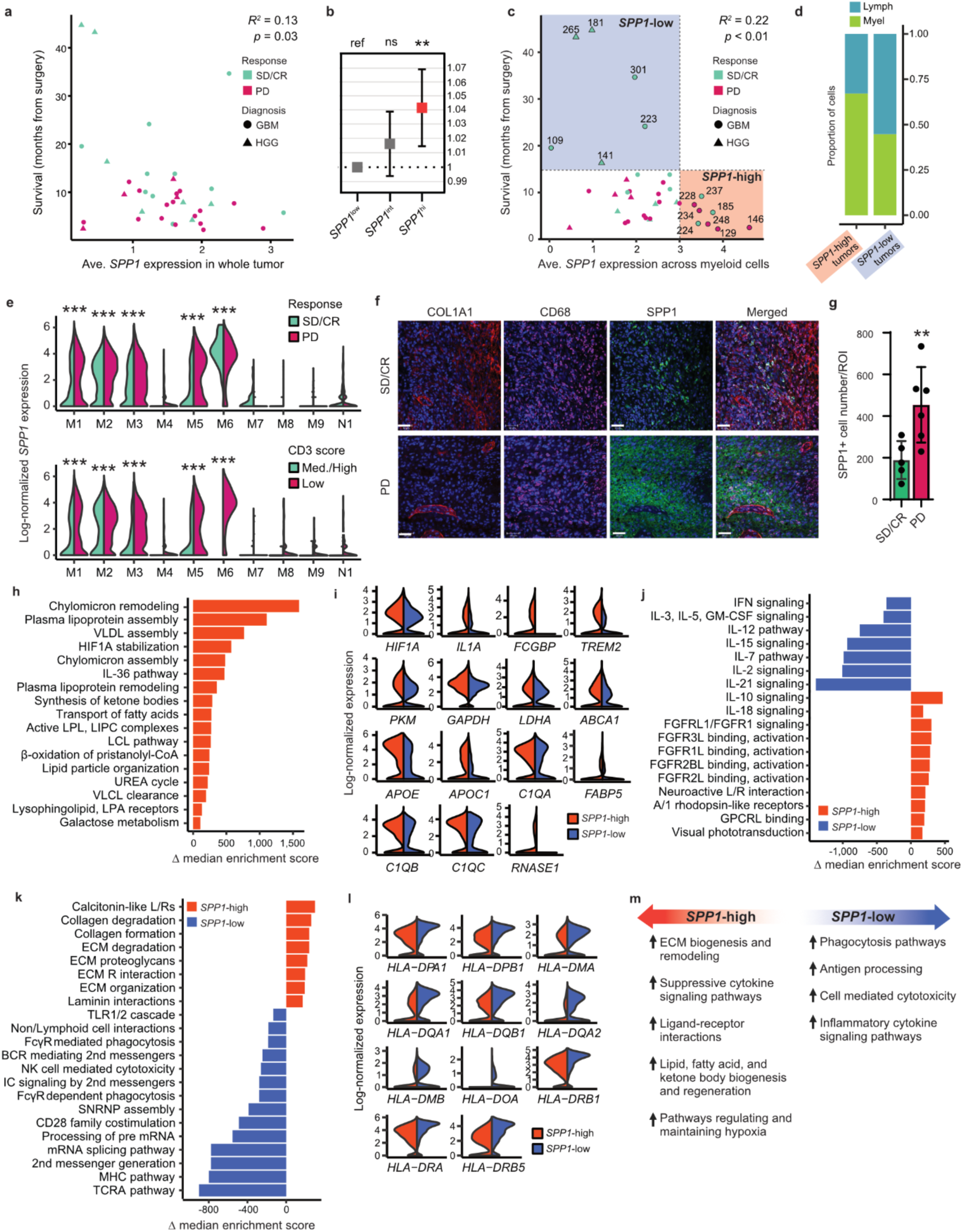
Characterization and comparative analysis of *SPP1* expressing myeloid cells and the association with resistance to IL13Rα2 CAR T cell therapy in GBM/HGGs. a, Expression of *SPP1* in GBM/HGG whole tumors is negatively correlated with survival (n = 37, response evaluable patients *R*^2^ = 0.13, p = 0.03). **b**, Cox’s proportional-hazard ratio (HR) for survival by the proportion (%) of myeloid cells expressing *SPP1* (*SPP1*^int/hi^) **a**mong 37 evaluable GBM/HGG patients. The HR = 1.04 (CI = 1.01–1.11, p = 0.002) indicating 4% increased risk with 1% increase in the relative abundance of *SPP1*^hi^ myeloid cells, is shown. **c**, *SPP1* expression in the myeloid fraction is significantly negatively correlated with survival (n = 37 response evaluable patients, *R*^2^ = 0.22, p = 0.01). Tumors representing the extremes in *SPP1* expression and survival post-surgery are annotated with their unique patient numbers (tumors encircled in blue designated “*SPP1*-low”, those encircled in orange designated “*SPP1-*high”). *R*^2^ is indicated. **d**, Differential proportions of myeloid and lymphoid cells in the immune cell fraction of *SPP1*-high and *SPP1*-low GBM/HGG tumors (*Χ*^2^ = 9.23 p-value = 0.002). **e**, *SPP1* expression (log-normalized) across myeloid subclusters in the scRNA-seq of SD/CR vs. PD tumors (top) or CD3 Med./High vs. CD3 Low tumors (bottom). *** denotes differential expression (two-sided Wilcoxon Rank Sum test, adj. p < 0.001) of *SPP1* in a given myeloid subcluster. **f**, Immunofluorescent staining for SPP1, COL1A1, and CD68 shows a higher expression of *SPP1* and enrichment of fibroblasts in PD tumors compared to SD/CR tumors. Bars indicate 50 µm. **g**, Mean ± SD. *SPP1*-expressing cell counts by immunofluorescent staining of tumor sections from patients that exhibited either SD/CR (n = 5) or PD (n = 6). ROI = 2 mm^2^, **, p = 0.02 using an unpaired t-test. **h**, Gene-set enrichment analysis reveals the significant upregulation of pathways involved in non-glycolytic ATP production in the myeloid fraction of *SPP1*-high tumors as compared to *SPP1*-low tumors. The delta of the median enrichment score is presented, with positive values indicating upregulation in *SPP1*-high. Significantly differentially regulated pathways were selected with a two-sided Wilcoxon FDR-p < 0.01, and a subset of significant pathways were selected for plotting. **i**, Expression (log-normalized) of canonical markers involved in non-glycolytic ATP production, differentially expressed in the myeloid fraction of GBM/HGGs representing the extremes denoted in panel (c). Differentially expressed genes with two-sided Wilcoxon Rank Sum test FDR-p < 0.001 when comparing *SPP1*-high vs. low are depicted. **j**, Significant downregulation of inflammatory pathways, upregulation of suppressive interleukin pathways, and higher receptor-ligand interactions (e.g., FGF receptors and associated ligands) in the myeloid fraction of *SPP1*-high tumors as compared to *SPP1*-low tumors. The delta of the median enrichment score is presented; significantly differentially regulated pathways were selected with a two-sided Wilcoxon FDR-p < 0.01, and a subset of significant pathways were selected for plotting. **k**, Upregulation of pathways involved in ECM synthesis and remodeling, and downregulation of pathways involved in antigen processing and presentation in the myeloid fraction of *SPP1*-high tumors as compared to *SPP1*-low tumors. The delta of the median enrichment score is presented; significantly differentially regulated pathways were selected with a two-sided Wilcoxon FDR-p < 0.01, and a subset of significant pathways were selected for plotting. **l**, Expression (log-normalized) of canonical markers involved in antigen presentation in the myeloid fraction of GBM/HGGs representing the extremes denoted in (c). Differentially expressed genes with two-sided Wilcoxon Rank Sum test p < 0.001 when comparing *SPP1*-high vs. low are depicted. **m**, Summary of differential functional pathways in the myeloid fraction of GBM/HGGs, representing the extremes denoted in (c).

By contrast, in non-responding patients (PD, CD3-low), extracellular matrix signaling pathways were significantly up-regulated (Fig. 1e; Extended Fig. 2a). This included increased intercellular signaling for extracellular matrix components such as collagen, Cyclophilin A (CyPA), and Visfatin (Fig. 1e, Extended Fig. 2a). Notably, the extracellular matrix protein SPP1 (Osteopontin; OPN) emerged as the most dominant intercellular communication pathway in non-responding patients (Fig. 1e, Extended Fig. 2a).

Heightened intercellular communication was observed within *SPP1*-enriched myeloid cells and between *SPP1*+ myeloid and fibroblast clusters (Fig. 1f, Extended Fig. 2d). Further analysis revealed that *SPP1*+ myeloid cells, specifically clusters M1, M2, M3, M5 and M6, as well as fibroblast cluster F3, show high expression of SPP1 and engage through three dominant cell-cell interaction pathways: *SPP1-CD44*, *MIF-CD44+CD74*, and *TREM2-APOE* (Fig. 1g-h, Extended Fig. 2d-e). These pathways are known to be associated with suppressive signaling within the TME^11–13^. While high expression levels of *SPP1* have been linked to poor prognosis in multiple solid tumors^14–16^ the role of *SPP1* in mediating resistance to CAR T cell therapies is noticeably understudied^17,18^.

### Multifunctional *SPP1*-expressing myeloid cells are associated with therapeutic resistance to CAR T cell therapy in GBM/HGGs

To further elucidate the clinical significance of *SPP1* expression in response to CAR T cell therapy, we first analyzed its relationship with survival across GBM/HGG tumors, revealing a significant inverse correlation (*R*^2^ = 0.13, p = 0.03; Fig. 2a). Given that *SPP1* was primarily derived from myeloid cells, we next examined whether the abundance of *SPP1*-high myeloid cells correlated with survival and found a similarly significant inverse relationship, indicating their role in driving CAR T cell resistance (Fig. 2b). Further analysis confirmed a negative correlation between *SPP1* expression in myeloid cells and survival in GBM/HGG patients (Fig. 2c, Extended Fig. 2f-i). *SPP1*-high tumors exhibited a shift toward myeloid dominance with a reduced lymphoid-to-myeloid ratio, further highlighting the role of *SPP1*-expressing myeloid cells in modulating therapeutic response (*Χ*^2^ = 9.23 p-value = 0.002, Fig. 2c, d).

We next sought to determine the association of *SPP1* expression with a suppressive niche linked to poor outcomes in GBM/HGGs, by evaluating differential expression of *SPP1* across myeloid subclusters (Fig. 2e, Supplementary Tables 7-8). Of the five myeloid subclusters identified to have high to intermediate *SPP1* expression (*SPP1*^hi/int^; M1, M2, M3, M5 and M6), significantly higher *SPP1* expression was observed in patients displaying poor outcomes to CAR T cell therapy, both PD and CD3-low tumors (Fig. 2e). Of note, the M6 subcluster was never present in CD3-high tumors and its abundance was significantly increased in PD patients (Extended Fig. 2b,c). M6 is marked by elevated expression of immunosuppressive genes such as *C1Q* subtypes, *CD163*, *CCL4*, *TYROBP*, *TREM2*, *APOE*, and is one of myeloid subclusters with the highest expression of *SPP1* (Fig. 1f)^19–23^. While M1, M2, M3, and M5 make up a significant portion of the suppressive myeloid cell population within the tumors and higher *SPP1* expression was associated with poor outcomes to CAR therapy, total subcluster abundance was not significantly differentially distributed in outcome comparisons (Fig. 1f, Extended Fig. 2b,c). Additionally, the fibroblast subcluster F3 exhibits high *SPP1* and *APOE* expression and is enriched in PD and CD3-low non-responding groups compared to their corresponding counterparts (Fig. 1f, Extended Fig. 2b,c). In contrast, M4, M8, and M9 are *SPP1*-negative myeloid cells lacking suppressive features, such as *TREM2*, *IL1B*, *NLRP3*, and *CCL4* (Fig. 1f, Extended Fig. 2b,c). Of note, M9 is a *CD4*+ subcluster with upregulation of *GZMB* potentially representing activated myeloid cells and is found in greater abundance in SD/CR group (Fig. 1f, Extended Fig. 2b,c). Immunofluorescent staining of PD tumor samples compared to SD/CR tumor samples confirmed differences in *SPP1* expression non-responding patients, consistent with transcriptomic findings (Fig. 2f-g). Collectively, these findings highlight a potential unrealized role for *SPP1* in suppressing the response to CAR T cell therapy.

Bridging transcriptomic data to the functional states of myeloid cells, we investigated the multifunctionality of the myeloid niche associated with CAR T cell resistance by comparing myeloid cells from *SPP1*-high patient tumors exhibiting poorer responses and *SPP1*-low tumors exhibiting better responses (Fig. 2c). To this end, we calculated gene set enrichment across curated pathways on a single cell level. *SPP1*-high myeloid cells demonstrated significant enrichment in lipid metabolism and oxidative phosphorylation pathways, characterized by upregulation of lipid-related genes such as *APOC1, APOE,* and *FABP5* (Fig. 2h-i, Supplementary Table 9). This metabolic profile suggests an increased reliance on non-glycolytic ATP production, indicative of their suppressive phenotype.

In addition to their altered metabolic profile, *SPP1*-high myeloid cells displayed upregulation of suppressive interleukin (IL) signaling pathways (Fig. 2j), in stark contrast to the inflammatory IL pathways enriched in SPP1-low myeloid cells. Elevated ligand-receptor interactions, particularly involving FGFR and its ligands, were also observed in *SPP1*-high myeloid cells (Fig. 2j). Furthermore, pathways associated with increased ECM activity and hypoxia, along with decreased phagocytosis and a marked downregulation of antigen processing and presentation pathways were associated with *SPP1*-high myeloid cells (Fig. 2k-l, Supplementary Tables 9-10). Together, these data demonstrate a coordinated phenotypic shift associated with increased *SPP1* expression consistent with driving a suppressive TME (Fig. 2m).

### IFN-deficient solid tumors suppress CAR T cell responses through an altered suppressive TME that is linked to increased *SPP1*+ macrophages

Based on the findings that *SPP1*-high myeloid cells in human GBM/HGGs were associated with depleted IFN signaling and antigen presentation, we performed parallel orthogonal studies in syngeneic mouse models to explore the dependency of CAR T cell responses on intact tumor IFN signaling and antigen presentation. While defective IFN signaling has been shown to impair CAR T cell-mediated killing via direct tumor cell-intrinsic mechanisms involving T cell-tumor cell interactions^24–26^, the broader impact on shaping the TME and influencing CAR T cell responses remains largely unexplored. To this end, we leveraged JAK1-knockout (JAK1/KO) melanoma tumors (YUMM), which are defective in type I and II IFN signaling and exhibit low baseline MHC expression and are known to be resistant to immune checkpoint blockade and adoptive transfer of TCR-targeted tumor-specific T cells due to defective antigen presentation^27–29^.

We next established IL13Rα2-engineered YUMM JAK1/KO and WT tumor sublines and verified altered CAR T cell-induced gene transcription in JAK1/KO YUMM line compared to the YUMM WT counterpart (Extended Fig. 3a-c). We subsequently established syngeneic mouse models which are targetable by a mouse equivalent to our human clinical IL13Rα2-targeted CAR (Extended Fig. 3d)^30^. Tumor cells were injected subcutaneously into immunocompetent mice and treated with murine IL13Rα2-targeted CAR (mIL13Rα2 CAR) T cells administered intravenously (IV; Fig. 3a). YUMM JAK1/KO tumors demonstrated significant resistance to IV delivered mIL13Rα2 CAR T cell therapy as compared to WT YUMM counterparts (Fig. 3b). Further, JAK1/KO tumors displayed a significantly reduced number of circulating CD3+ and CAR T cells in the blood (Fig. 3c,d). Similar results were also observed when CAR T cells were administered intratumorally (IT) (Fig. 3e,f), suggesting that the limited therapeutic activity in JAK1/KO tumors could not solely be explained by defects in trafficking. Prior studies have shown that IFN signaling deficient tumors are resistant to CAR T cells due to deficits in upregulation of adhesion molecules in response to IFNγ, such as ICAM1, which can modulate optimal CAR T cell-tumor synapse formation and subsequent tumor cell killing. We therefore engineered JAK1/KO YUMM tumors to overexpress ICAM1; however, resistance to CAR T cell therapy persisted despite ICAM1 overexpression (Extended Fig. 4). Further *in vitro* cytotoxicity assays revealed comparable CAR T cell-mediated killing between JAK1/KO and WT YUMM cells, suggesting that intrinsic tumor susceptibility to CAR T cells remained intact (Extended Fig. 5).

**Figure 3.**
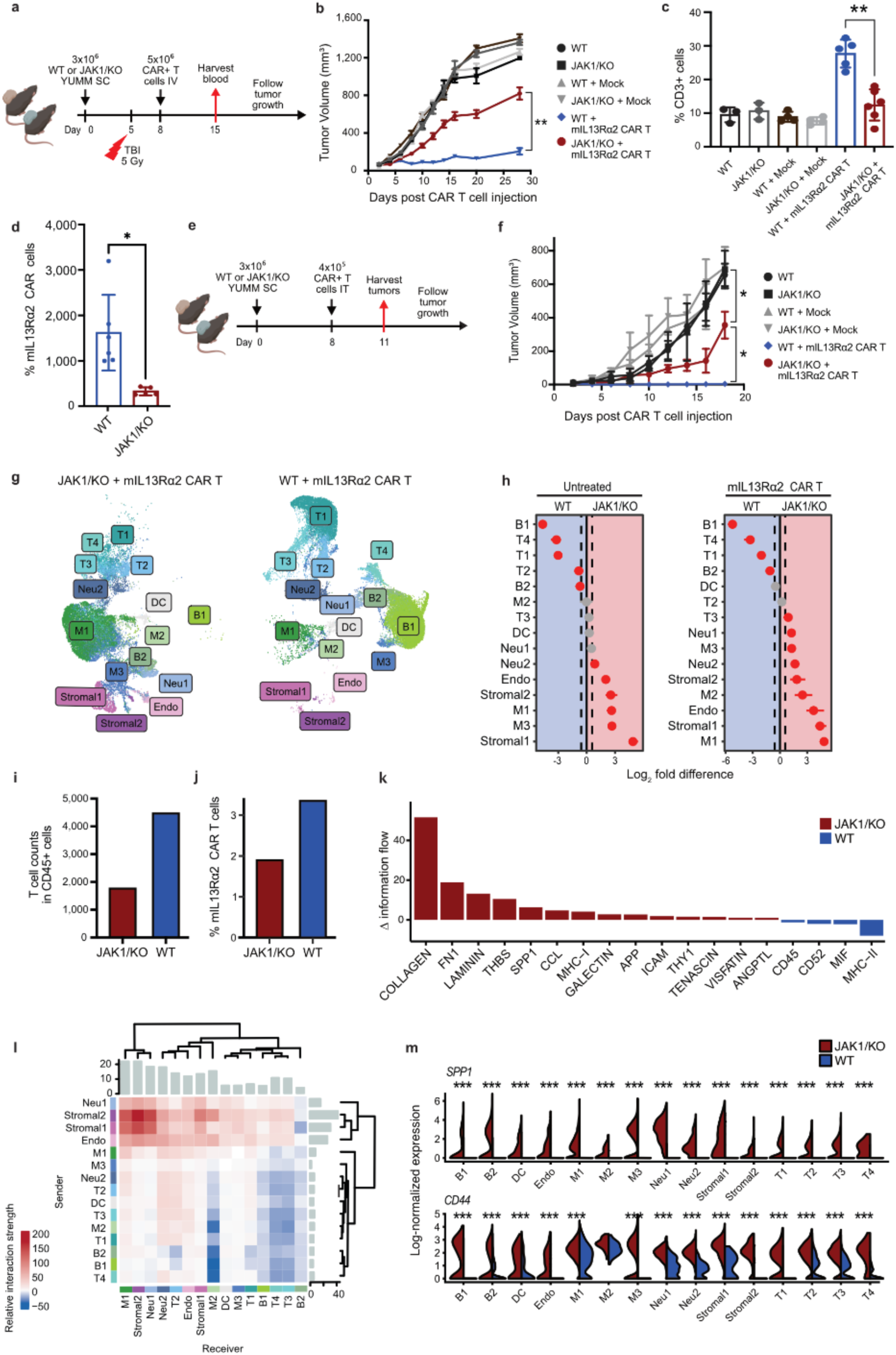
Characterization of IFN signaling-deficient tumor mouse model displaying resistance to mIL13Rα2 CAR T cell therapy. **a**, Schematic illustration of the study design for in vivo evaluation of systemic mIL13Rα2 CAR T cell therapy (IV) against YUMM WT (n = 5) vs. JAK1/KO (n = 5) tumor bearing mice. **b**, Differential tumor growth dynamics of YUMM JAK1/KO tumors compared to YUMM WT to systemic mIL13Rα2 CAR T cell therapy (IV). Significant differential tumor growth is denoted with asterisks (** p<0.01). **c**, Flow cytometric analysis shows a significantly reduced percentage of CD3+ cells in the blood of mice bearing YUMM JAK1/KO tumors treated with mIL13Rα2 CAR T cells compared to YUMM WT (** p<0.01). Data are mean ± SD. **d**, Flow cytometric analysis showing reduced CAR T cell percentages in the blood of YUMM JAK1/KO tumor-bearing mice compared to YUMM WT (* p<0.05). Data are mean ± SD. **e**, Schematic illustration of the study design for *in vivo* evaluation of intra-tumoral mIL13Rα2 CAR T cell therapy against WT (n=7) vs. JAK1/KO (n=7) YUMM tumor bearing mice. **f**, Volumes of tumors untreated or treated with mIL13Rα2 CAR T cells, exhibiting significant response in YUMM WT tumors in comparison to suboptimal response in YUMM JAK1/KO tumors (* p<0.05). Data are mean ± SEM. **g**, Clustering of scRNA-seq data of 51,820 CD45+ cells from 6 JAK1/KO (23,848 cells, including 14,831 cells from mIL13Rα2 CAR T treated and 9,017 cells from untreated control tumors) and 6 WT (27,972 cells, 14,128 treated, 13,844 untreated) tumors, annotated by cluster identity. **h**, Differences in cell type distribution between YUMM JAK1/KO and YUMM WT tumors untreated (left) or treated with mIL13Rα2 CAR T cell therapy (right). Log_2_ fold-differences with 95% confidence intervals are presented. Results for cell types with significant differences based on a permutation p-value are highlighted in red. **i**, T cell counts in scRNA-seq data reveal reduced infiltration of CD3+ T cells in JAK1/KO YUMM tumors compared to WT counterparts following mIL13Rα2 CAR T cell treatment. **j**, The percentage of mIL13Rα2 CAR T cells identified in scRNA-seq analysis shows a reduction in JAK1/KO YUMM tumors relative to WT YUMM tumors. **k**, Differentially regulated ligand-receptor signaling pathways in a comparison of the TME of CAR treated YUMM JAK1/KO and WT tumors. Significant pathways were selected with a p<0.01, and pathways with an absolute delta of >0.8 were selected for plotting. The information flow for a given signaling pathway is defined by the sum of communication probability among all pairs of cell groups in the inferred network. Delta of the overall information flow between groups in each comparison is presented, with positive values indicating increased signaling in JAK1/KO. **l**, Differential strength of interactions between all pairs of analyzed cell types in a comparison of CAR treated JAK1/KO and WT tumors. Positive values indicate increased signaling in JAK1/KO. **m**, *SPP1* and *CD44* are upregulated across TME subclusters of YUMM JAK1/KO tumors compared to YUMM WT tumors after mIL13Rα2 CAR T cell therapy. Log-normalized *SPP1* expression is stratified by JAK1 and treatment status. Significant differential expression is denoted with asterisks: * p<0.05, ** p<0.01, *** p<0.001.

**Figure 4.**
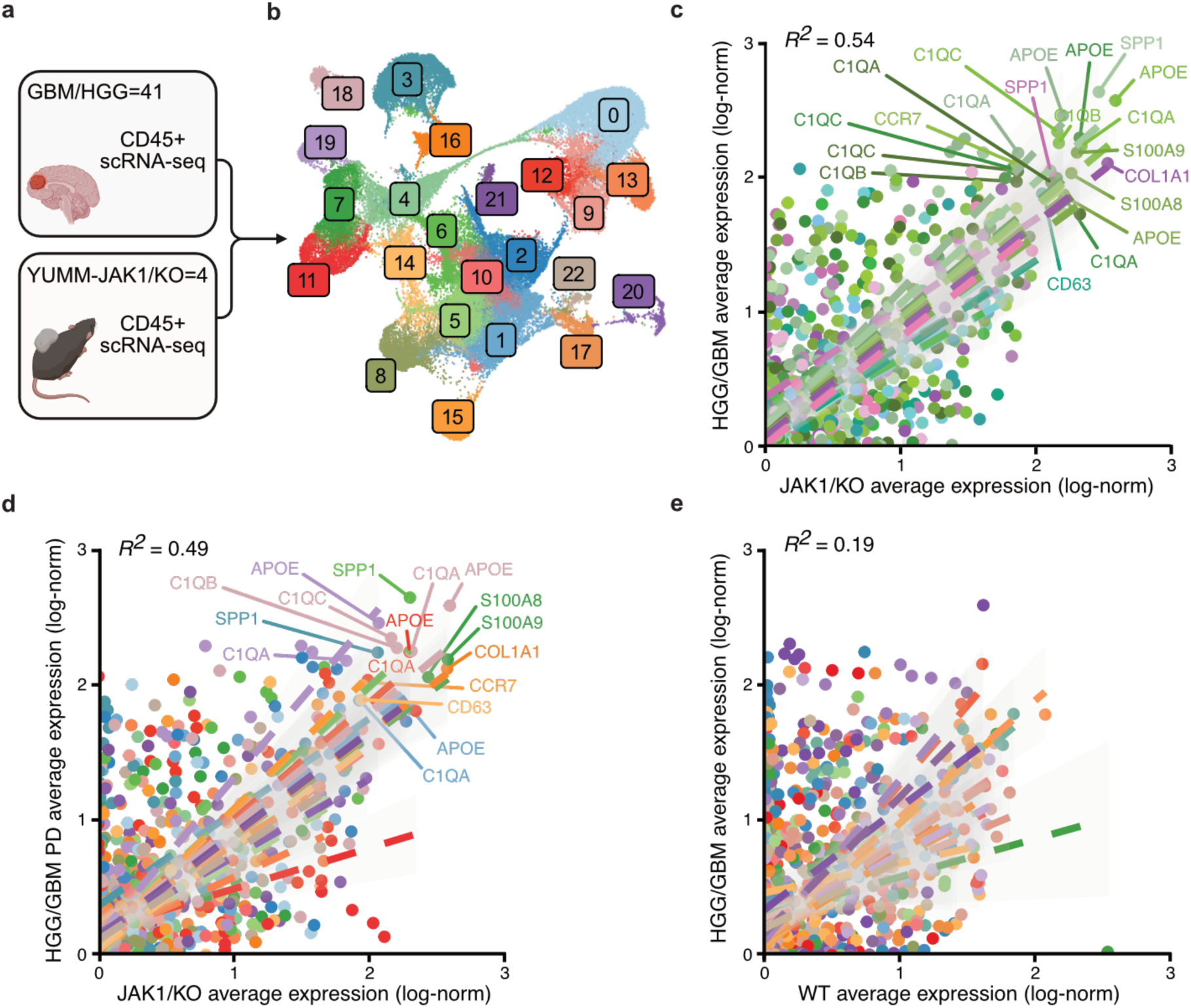
Concordant suppressive programs in human GBM/HGG and mouse IFN signaling-deficient tumors. **a**, Schematic illustration of the integrated analysis of scRNA-seq data from patient GBM/HGG and mouse YUMM JAK1/KO tumors. Immune and stromal cells from human GBM/HGG (n = 41) and YUMM JAK1/KO mouse melanoma tumors (n = 4) were jointly analyzed to examine regulatory programs shared between the two tumor types. **b**, UMAP dimensionality reduction of GBM/HGG (28,156 cells) and YUMM JAK1/KO (40,324 cells) scRNA-seq data, annotated by cluster identity. **c**, Expression of canonical myeloid and fibroblast markers (presented in Fig. 1f) are highly correlated (*R*^2^ = 0.55) between GBM/HGG and JAK1/KO YUMM tumors in relevant cell types. Colors of cell types correspond to Fig. 1f. Subsets of highly expressed genes are notated. Regression lines with confidence intervals are depicted for each gene–cell type combination. **d**, Expression of selected markers (as in panel c) between the TME subsets of GBM/HGG tumors with best response of Progressive Disease and JAK1/KO YUMM tumors (*R*^2^ = 0.41). **e**, Expression of selected markers (as in panel c) is poorly correlated (*R*^2^ = 0.04) between the TME subsets of GBM/HGG and JAK1/WT YUMM tumors.

**Figure 5.**
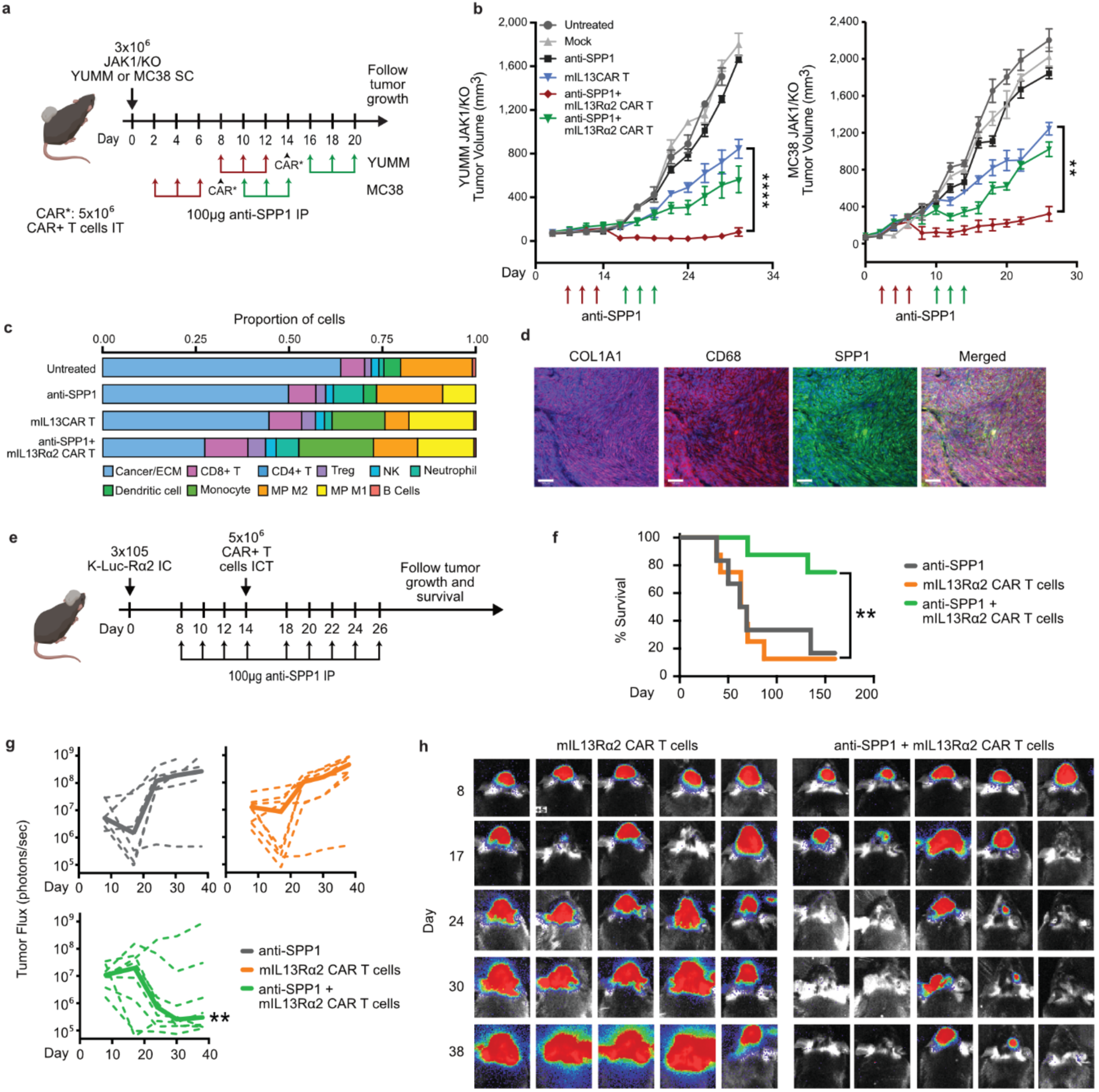
Blocking SPP1 in the tumor microenvironment enhances mIL13Rα2 CAR T cell therapy against IFN signaling-deficient and glioma tumors in syngeneic mouse models. **a**, Schematic illustration demonstrating experimental design combining anti-SPP1 antibody and mIL13Rα2 CAR T cell therapy against YUMM JAK1/KO (n=5 per arm) and MC38 JAK1/KO tumors (n=5 per arm). **b**, Tumor volumes comparing combination therapy to mIL13Rα2 CAR T cell therapy alone and anti-SPP1 antibody treatment alone in YUMM JAK1/KO and MC38 JAK1/KO tumors. ** p<0.01, **** p<0.0001. Data are mean ± SEM. **c**, Cellular deconvolution of bulk RNA-seq from YUMM JAK1/KO tumors at day 17 (72 hours post-mIL13Rα2 CAR T cell administration on day 14), comparing tumors treated with a combination of anti-SPP1 and mIL13Rα2 CAR T cells, mIL13Rα2 CAR T cells alone, anti-SPP1 antibody alone, and an untreated control group. **d**, Representative immunofluorescent staining of CD68, COL1A1, and SPP1 in KLuc orthotopic glioma tumors (Bars indicate 50 µm). **e**, Schematic illustration of experimental design of KLuc orthotopic glioma model treated with combination of anti-SPP1 antibody and mIL13Rα2 CAR T cell therapy compared to mIL13Rα2 CAR T cell therapy alone and anti-SPP1 antibody alone groups (n=5 per arm). **f**, Survival analysis revealed significantly longer survival of KLuc harboring mice treated with combination therapy compared to mIL13Rα2 CAR T alone, anti-SPP1 antibody alone treated groups. Data are mean ± SEM. **g**, Flux values revealed significantly lower bioluminescence values in KLuc harboring mice treated with combination therapy compared to mIL13Rα2 CAR T alone, anti-SPP1 antibody alone treated groups. **h**, Mice heads showing the flux values comparing mIL13Rα2 CAR T and anti-SPP1 antibody combined treated group to mIL13Rα2 CAR T alone group.

We next evaluated the potential role of the TME in mediating resistance to mIL13Rα2 CAR T cells in JAK1/KO models. We performed paired single cell analyses of YUMM JAK1/KO and WT tumors following mIL13Rα2 CAR T cell IT treatment (Fig. 3g, Extended Fig. 6a, Supplementary Table 13). We found cell type differences between WT and YUMM JAK1/KO were largely consistent in both untreated and mIL13Rα2 CAR T treated mice, suggesting the endogenous suppressive TME is maintained through therapy (Fig. 3h, Extended Fig. 6a). We found a significant enrichment of macrophages and stromal cells in the YUMM JAK1/KO mice and an enrichment of B and T cells in the WT mice (Fig. 3h-i). Furthermore, from our scRNA-seq analysis we were able to identify the proportion of mIL13Rα2 CAR T cells in the tumors and found a significantly lower proportion within YUMM JAK1/KO compared to WT (Fig. 3j). Interestingly, we see higher expression of *CD44* and *PD1* expression in mIL13Rα2 CAR T cells in YUMM JAK1/KO compared to WT counterparts (Extended Fig. 6).

Comparative cell-cell signaling analyses highlighted significant differences for intercellular communication pathways in YUMM JAK1/KO tumors, particularly those regulating ECM scaffold, remodeling and regeneration, such as collagen, fibronectin, and laminin (Fig. 3k). Consistent with findings from GBM/HGGs, *SPP1* emerged as one of the top upregulated pathways in YUMM JAK1/KO tumors, while *MHCII* was notably upregulated in WT tumors (Fig. 3k, Fig. 1e). Cell-cell communication analysis further demonstrated increased interaction probabilities between fibroblasts and myeloid cells in YUMM JAK1/KO tumors relative to YUMM WT, with *SPP1* representing one of the most prominent pathways driving these interactions (Fig. 3k-l). Notably, both *SPP1* and its cognate receptor *CD44* were significantly upregulated across all immune-driven populations, including fibroblasts, neutrophils, and myeloid cells, in JAK1/KO tumors compared to YUMM WT (Fig. 3m).

Concurrent Cytometry by Time-of-flight analysis (CyTOF) analysis corroborated the transcriptomic findings, revealing elevated populations of suppressive macrophages, fibroblasts, exhausted T cells, and follicular T cells, along with a pronounced reduction in activated and naive T cells in YUMM JAK1/KO tumors compared to YUMM WT tumors post mIL13Rα2 CAR T therapy (Supplementary Table 14, Extended Fig. 7b-e). Immunofluorescence staining further confirmed significantly enhanced *SPP1* expression in YUMM JAK1/KO tumors compared to WT following mIL13Rα2 CAR T cell therapy (Extended Fig. 8). Together, these data demonstrate how the IFN signaling-deficient JAK1/KO tumor microenvironment fosters an immunosuppressive niche, markedly involving *SPP1* and contributing to resistance against mIL13Rα2 CAR T therapy.

### Conserved TME features associated CAR T cell resistance across solid tumors

We next conducted an integrative analysis comparing scRNA-seq data from GBM/HGG patients paired with mouse JAK1/KO and WT YUMM tumors (Fig. 4a,b). Strikingly, this analysis revealed a high concordance between GBM/HGGs and IFN-deficient, but not WT, mouse melanoma tumors, highlighting the universality and conservation of key TME features driving resistance to CAR T cell therapy in solid tumors (Fig. 4c-d). GBM/HGGs and JAK1/KO tumors showed strong concordance in the high expression of canonical suppressive macrophage and fibroblast markers. These genes include *APOE*, *C1QA*, *C1QB*, *C1QC*, *COL1A1*, and *SPP1*, all expressed in multiple integrated macrophage subclusters, as well as *S100A8* and *S100A9*, which were expressed in neutrophil subclusters. In contrast, these genes were not as correlated in the comparison of GBM/HGGs and YUMM WT tumors that were more responsive to CAR T cell therapy (Fig. 4e). Collectively, the expression of these conserved signatures across immune components predominantly shapes the suppressive TME, thereby reinforcing its unresponsiveness to CAR T cell therapy. These findings suggest that despite diverse tumor-intrinsic resistance mechanisms to CAR T cell therapy—such as those observed in patient GBM/HGG and IFN signaling-deficient mouse tumors—resistance ultimately is conferred by shared suppressive features of the TME, primarily driven by *SPP1*+ myeloid populations and extracellular matrix remodeling.

### SPP1 blockade overcomes CAR T resistance in solid tumors

Given the convergence of SPP1 mediated signaling associated with an immuno-suppressive TME, we hypothesized targeting SPP1 alongside CAR T cell therapy may overcome resistance. To test this, we used three preclinical models of solid tumors with known suppressive TMEs and IFN deficiencies. First, we returned to the YUMM JAK1/KO melanoma model used in the studies above as well as extending to an additional IFN signaling deficient tumor model, MC38 JAK1/KO (Fig. 5a,b). MC38 JAK1/KO shows resistance to checkpoint blockade therapy and recapitulates many of the immunosuppressive features seen in the YUMM JAK1/KO^27^. Similar to YUMM tumor sublines, MC38 JAK1/KO is engineered to express IL13Rα2. In both models we employed a dual approach using an anti-SPP1 blocking antibody in combination with mIL13Rα2 CAR T cell therapy. In one arm, we administered three doses of anti-SPP1 antibody prior to mIL13Rα2 CAR T cell therapy, while in another arm, anti-SPP1 was delivered concurrently with mIL13Rα2 CAR T cells and continued for three doses (Fig. 5a). Analysis of tumor volumes revealed a significant reduction in the size of tumors in the sequentially treated group compared to a modest reduction in the concomitantly treated group (Fig. 5b-left panel). Similar results were observed in MC38 JAK1/KO models, where pre-treatment with anti-SPP1 followed by mIL13Rα2 CAR T cell therapy demonstrated significant tumor responses (Fig. 5b-right panel). RNA bulk sequencing of YUMM JAK1/KO tumors primed with anti-SPP1 before CAR T cell therapy revealed increased levels of dendritic cells, activated T cells, and inflammatory macrophages compared to tumors treated with anti-SPP1 or CAR T cells alone (Fig. 5c).

To further demonstrate the impact of SPP1 modulation on CAR T therapy, we extended these studies to a mouse glioma model. We stained tissues from three distinct syngeneic glioma tumors for SPP1, revealing elevated expression across all tumors, with particularly high levels in Kluc tumors (Fig. 5d, Extended Fig. 8). Kluc is a firefly luciferase-engineered subline of KR158 derived from spontaneous gliomas in Nf1, Trp53 mutant mice, mimics the invasive features of GBM/HGG and demonstrates notable resistance to immunotherapies. Furthermore, tissue from Kluc tumors recapitulate SPP1 driven features associated with poor CAR T outcomes in human GBM/HGG (Fig. 5d, Fig. 2f). As above, we combined anti-SPP1 therapy with CAR T cell therapy against Kluc orthotopic tumors. Both survival analysis and tumor flux values revealed that the mice primed with anti-SPP1 and continuing to receive anti-SPP1 after CAR T cell therapy exhibited significantly improved outcomes compared to tumors treated with anti-SPP1 alone or CAR T cells alone (Fig. 5e-h). Together, these preclinical findings support the notion that SPP1 fuels the suppressive niche and promotes tumor resistance to CAR T cell therapy. Sequential blockade of SPP1 and mIL13Rα2 CAR T cell therapy overcomes resistance in multiple tumor models.

## Discussion

Despite a notable recent surge in reported trials using CAR T cells against solid tumors, their efficacy is still limited^31,32^. Recent advances have shed light on the role of TME in immunotherapy resistance within solid tumor^33–35^, yet the complex interplay of intercellular signaling and macrophages, crucial components of the suppressive TME, remains largely unexplored in the context of response to CAR T cell therapy. These complexities pose challenges in unraveling the dynamics of resistance to CAR T cell therapies amidst the intricacies of patient-specific TMEs.

Through a clinically oriented single-cell analysis of patients with gliomas undergoing IL13Rα2-targeted CAR T cell therapy, we examined the distribution of TME cellular constituents, uncovering differences in the abundance of macrophages and the activation of their distinct extracellular signaling networks in responsive versus non-responsive tumors. Notably, *SPP1*-expressing macrophages emerged as a pivotal determinant of response to CAR T cell therapy, while other examined extracellular pathways did not exhibit such a strong relevance to treatment outcome. Our subsequent profiling of high *SPP1*-expressing macrophages unveiled distinct pathways that differentiate them from other macrophage subsets. These macrophages exhibited unique metabolic and functional adaptations, including upregulated lipid metabolism, enhanced receptor-ligand interactions—many of which remain largely unexplored—elevated ECM biogenesis and remodeling, and suppressive interleukin signaling pathways. Conversely, they demonstrated reduced antigen presentation and phagocytic activity, collectively fostering a suppressive TME and driving resistance to therapy. Further research is essential to mechanistically investigate the factors contributing to increased *SPP1* expression in nonresponsive tumors.

The interdependent tumor-TME ecosystem is critically regulated by IFN signaling, which plays a pivotal role in mediating antigen presentation and governing macrophage antitumor plasticity^36–38^. Moreover, CAR-T cell mediated IFN-production is critical for productive antitumor responses, playing an essential role in enhancing antigen presentation and reshaping the TME^30,38^. In marked contrast, hypoxic conditions, coupled with the deprivation of IFN signaling, result in impaired antigen presentation and reprogram macrophages toward SPP1-dominated suppressive states^38,39,40^. Disruptions in IFN signaling pathways, such as reduced tumor IRF8 activity, intensify *SPP1*-mediated immunosuppression and *CD44*-dependent suppression of T cell function^41,42^. These findings highlight the critical role of IFN signaling in orchestrating immune dynamics and shaping TME responsiveness to immunotherapy^43^. In some contexts, IFN-deficient tumors display resistance to CAR T cell killing *in vitro*, due to lower adhesion molecule expression and suboptimal CAR-tumor synapse formation^24,25^. By contrast, we show that IL13Rα2 CAR T cells have comparable *in vitro* killing potency against both WT and JAK1/KO IFN-deficient tumors, likely due to the high affinity of this IL13-ligand CAR for its receptor^44^, which may compensate for IFN-dependent effects on tumor binding and avidity. Instead, the stark resistance of IFN-deficient tumors to CAR T cell therapy *in vivo* is predominantly attributed to TME-driven effects, with JAK1 knockout tumors exhibiting significantly altered immune cell expression compared to their wild-type counterparts.

Integrative single-cell RNA sequencing analysis of patient gliomas and an IFN-signaling deficient mouse melanoma model allowed us to take a novel approach to dissecting the TME as a driver of solid tumor resistance to CAR T cell therapy. Consistent patterns identified in the signatures of suppressive macrophages, along with the orchestrated intercellular signaling networks facilitating suppressive interactions within the TME, suggest shared mechanisms driving CAR T cell resistance across diverse solid tumor types. These mechanisms may potentially be co-regulated by distinctly different intracellular pathways, warranting further investigation. The prevalent abundance and distinctive signature of *SPP1* in non-responsive versus responsive solid tumors to CAR T cells, evidenced by a single-cell level analysis of multiple data modalities and functional phenotyping, together with translational outcomes targeting SPP1 alongside CAR T cell therapy in syngeneic mouse models, indicate a promising avenue for altering the TME landscape and advancing current therapeutic strategies. However, considering the adaptable nature of the TME under CAR T cell-associated immunomodulation and the subsequent emergence of other suppressive niche constituents, it is imperative to mechanistically investigate the factors leading to *SPP1* upregulation and macrophage cellular states resulting in increased expression of *SPP1* molecules.

While previous studies have delineated TAM polarity through a binary framework such as *CXCL9* and *SPP1* or the M1 and M2 paradigm^17^, our approach emphasized the concept of immune cell multipolarity, with *SPP1* emerging as a pivotal identifier. Our analysis reveals that although universal suppressive markers, including *C1Qs, IL1B, APOE, CD163,* and *CD206,* consistently characterize and define a diverse spectrum of suppressive macrophages, *SPP1* itself is distinctively linked to clinical outcomes following CAR T cell therapy. Furthermore, our demonstration that SPP1-blockade reverses this immunosuppression and enhances CAR T cell antitumor responses underscores its unique role in modulating therapeutic efficacy. In *SPP1*-negative macrophages, the expression of other key markers, such as *PD1*, *PDL1*, and *CD4*, critically determines macrophage functional fate. Moreover, an in-depth examination of *SPP1*-positive and *SPP1*-negative macrophages uncovered fundamentally divergent functional transcriptomic profiles between these subsets. Thus, while universal suppressive markers broadly delineate the suppressive phenotype, *SPP1* distinguishes itself through its strong association with therapeutic outcomes, highlighting its singular significance in influencing responses to CAR T cell therapy. Extending observations from heterogeneous GBM/HGGs to peripheral solid tumors reveal conserved TME characteristics underlying resistance to CAR T cell therapy. These parallels underpinned universally dominant mechanisms driving therapeutic resistance, offering not only critical insights into immune evasion but also actionable strategies to mitigate resistance.

## Supplementary Tables

**Supplementary Table 1.** Numbers for GBM/HGG cells sequenced by batch and unique patient number (UPN).

**Supplementary Table 2.** Top markers for each cell type in the GBM/HGG scRNA-seq data.

**Supplementary Table 3.** Top markers for each cell type in the immune and stromal compartments of the GBM/HGG scRNA-seq data.

**Supplementary Table 4.** Numbers of cells by UPN and cell type.

**Supplementary Table 5.** Numbers of immune cells and fibroblasts by UPN and cell type.

**Supplementary Table 6.** CellChat communication probabilities for all significantly differentially activated ligand-receptor pairs in significant cell types.

**Supplementary Table 7.** Differentially expressed genes (adj. p <0.01, absolute log_2_FC >2) between CR/SD and PD tumors for each immune/fibroblast cell type.

**Supplementary Table 8.** Differentially expressed genes (adj. p <0.01, absolute log_2_FC >2) between CD3 high vs. low tumors for each immune/fibroblast subcluster.

**Supplementary Table 9.** Differentially expressed genes (adj. p <0.01, absolute log_2_FC >2) in a comparison of myeloid cells from tumors with upregulated SPP1 and shorter survival after surgery, vs. those with low *SPP1* expression and longer survival.

**Supplementary Table 10.** Differentially activated pathways (FDR-p<0.01) in a comparison of myeloid cells from tumors with upregulated *SPP1* and shorter survival after surgery, vs. those with low *SPP1* expression and longer survival.

**Supplementary Table 11.** Differentially activated pathways (FDR-p<0.01) in a comparison of myeloid cells from CR/SD and PD tumors.

**Supplementary Table 12.** Differentially activated pathways (FDR-p<0.01) in a comparison of myeloid cells from tumors with high (3-4) vs. low CD3 (0-2) scores.

**Supplementary Table 13.** Top markers for each cell type in the CD45+ YUMM tumor scRNA-seq data.

**Supplementary Table 14.** Panel of immune markers used in the Cytometry of Time of Flight (CyTOF) analysis.

**Supplementary Table 15.** Differentially activated pathways (FDR-p<0.01) in a comparison of myeloid cells from JAK1/KO vs. WT YUMM tumors.

## Author Contributions

SG and CEB conceived of this project. SG, HMN, and CEB wrote the primary draft of the manuscript and prepared figures. SG, HMN, NEB and CEB reviewed and edited the manuscript. SG led, designed, conducted and analyzed all experiments associated with this study. SX, SS, and CM designed, conducted and analyzed the *in vivo* and *in vitro* experiments. HMN and YZ performed the bioinformatics analysis of sc-RNAseq data. MA performed immunofluorescent staining on mouse and human samples. BCA performed and analyzed CyTOF studies. RAW, RS, BA, LP, MC and EDM provided technical assistance. DYT, AK and AR provided JAK1/KO cell lines. AK, DA, AR, SJF, BB, provided advisement and project input. SJF, BB, XW provided supervision of project areas. NEB and CEB supervised the overall study.

## Acknowledgments

This project was supported by California Institute for Regenerative Medicine (CLIN2-10248, C.E.B), Gateway for Cancer Research (G-14-600, C.E.B, B.B.), the Food and Drug Administration (R01FD005129, C.E.B, B.B), the National Institute of Health (R01 CA236500, C.E.B, B.B; R01 CA254271, C.E.B, B.B), the Mann Foundation (C.E.B, B.B), the Heritage Provider Network Foundation (C.E.B, B.B), the Norris Foundation, the Ivy Foundation, the Parker Institute for Cancer Immunotherapy (S.G.), and the Brenn Foundation (S.G.). We thank J.O. for contributions to the final figure and draft format.

## Methods

### GBM/HGG tumor samples acquisition and processing

Baseline tumor tissue samples were obtained from 41 patients with recurrent/refractory malignant glioma participating in a Phase I study (IRB #13384) on cellular immunotherapy using central memory-enriched IL13Rα2-targeting Chimeric Antigen Receptor (CAR) T cells. Tumor resection material was collected through the COH Department of Pathology according to the clinical protocol.

### GBM/HGGs library preparation and single-cell sequencing

Single-cell sequencing of cryopreserved freshly dispersed tumor samples from 41 individuals was carried out using the 10x Chromium platform. For Batch 1, patient PBMC’s exomes were collected via Illumina exome panel and sequenced at 20M read pairs per patient for sample deconvolution. For batches 2-6 and batches 42-1, 42-2, 43-1, 43-2, 44-2, 45, 46, and 47, BioLegend Total Seq-C hashtag antibodies were used to allow sample deconvolution after pooled processing, with barcoded samples sorted at equal proportions into a single collection tube. Samples in the remaining batches were sequenced without pooling. 60,000 cells were loaded to a single Gel Bead-in-Emulsion (GEM) reaction onto the Chromium instrument. ScRNA-seq library preparation was performed according to manufacturer protocols and sequenced on Illumina iSeq100 for cell count validation and NovaSeq6000 at the recommended depth. Sequence data were processed using 10x Genomics Cell Ranger V5.0 and Ensemble 98.

### GBM/HGG tumor single-cell data processing and analysis

Single-cell sequencing data were analyzed using Seurat v5^1^. CellRanger objects for each batch were imported to create a Seurat object for each of the 37 batches. For Batch 1, sample identities were deconvoluted, and multiplets were identified using Demuxlet. Exome sequencing FASTQ reads were chunked to 40M reads, aligned to GRCh38 using BWA, and processed with samtools (v1.10) fixmates and samtools sort (v1.10)^2^. Individual chunks were merged, and PCR and optical duplicates were marked with samtools. Genotypes were called with DeepVariant (https://github.com/google/deepvariant). For the batches with hash-antibodies, demultiplexing was done using the HTODemux() function of Seurat. The single-cell data were filtered to retain singlets with >500 unique RNA features detected, >1,000 RNA feature counts, and <10% of reads mapping to mitochondrial genes. After demultiplexing and quality filtering, we filtered the data for ambient RNA contamination using SoupX v1.6.2^3^. In the batches sequenced without multiplexing, possible doublets were identified using DoubletFinder v2.0.3^4^.

Gene expression data were normalized with SCT^5^, and samples were integrated using rPCA. The number of significant principal components (PCs) to be included in dimensionality reduction and unsupervised clustering was determined by calculating the difference between the proportion of variation associated with each PC and their subsequent PC and selecting the last point where the difference is more than 0.1%. Uniform Manifold Approximation and Projection (UMAP) was used for dimensionality reduction and 2D visualization of cell clusters.

The scRNA-seq data were analyzed for copy number alterations using the R package InferCNV v 1.20.0 (https://github.com/broadinstitute/inferCNV) with the following parameters: HMM_type = “i3, hclust_method = ’ward.D2’, sd_amplifier = 3, noise_logistic = T, tumor_subcluster_partition_method = ’random_trees’.

Highly expressed marker features for each cluster were identified using the presto (v1.0.0) implementation of the Wilcoxon rank test (https://github.com/immunogenomics/presto). Clusters were annotated based on cell type marker expression. Two clusters that did not express reasonable levels of any cell type markers and consisted mostly of cells from single individuals were excluded from downstream analyses. For the differential expression analysis between groups, we employed the Seurat implementation of the Wilcoxon rank sum test. We tested for the statistical significance of differences in cell type abundance between groups using the R package scProportionTest^6^.

Ligand-receptor interaction analysis was carried out using CellChat v2.1.1^7^ leveraging the CellChatDB human database (v1.0.0), which contains 1,939 validated molecular interactions, including 61.8% of paracrine/autocrine signaling interactions, 21.7% of extracellular matrix (ECM)-receptor interactions and 16.5% of cell-cell contact interactions. Expression data were preprocessed for the cell-cell communication analysis by identifying over-expressed ligands or receptors in one group and then identifying over-expressed ligand-receptor interactions. Then, gene expression data were projected onto the protein-protein interaction network using the function projectData(). Next, communication probabilities were computed using computeCommunProb(), and the results were filtered to retain instances with a minimum of ten cells in each group. Cell-cell communication was inferred at a pathway level by summarizing the communication probabilities of all ligands-receptor interactions associated with each signaling pathway using computeCommunProbPathway(), and the aggregated communication network was calculated by counting the number of links and summarizing the communication probability using aggregateNet().

To test the effects of cell type abundance and gene expression levels on overall survival, we employed Cox proportional-hazards model using the coxph() function of the R package survival v3.5-5 (https://github.com/therneau/survival). In survival analyses, data from 37 evaluable patients were used.

We used the R package escape v2.1.3^8^ to calculate enrichment scores for KEGG, REACTOME, BIOCARTA, and HALLMARK gene sets for each single cell. The enrichment calculation was carried out using the enrichIt() function, which utilizes the GSVA R package (v1.53.22) and the Poisson distribution for RNA.

### JAK1KO/WT tumor library preparation and single-cell sequencing

The mouse scRNA-seq library preparation was performed according to the manufacturer protocols and sequenced on Illumina iSeq100 for cell count validation and NovaSeq6000 at the recommended depth.

### JAK1KO/WT tumor single-cell data processing and analysis

Raw base call files of scRNA-seq data were analyzed using CellRanger (v5.0). The “mkfastq” command was used to generate FASTQ files and the “count” command was used to generate raw gene-barcode matrices aligned to the 10X Genomics GRCm38 reference genome (mm10). The data from all samples were combined in R (4.0.4) using the Read10X() function from the Seurat package (v4.0.3)^9^ and an aggregate Seurat object was generated. Filtering was conducted by retaining cells that had unique molecular identifiers (UMIs) greater than 400, expressed 200 and 9000 genes inclusive, and had mitochondrial content less than 15 percent. No sample batch correction was performed. Data were normalized using the “LogNormalize” method and using a scale factor of 10,000. Using Seurat’s ScaleData() function and “vars.to.regress” option UMI’s and percent mitochondrial content were used to regress out unwanted sources of variation. The number of variably expressed genes were calculated using the following criteria: normalized expression between 0.125 and 3, and a quantile-normalized variance exceeding 0.5. To reduce dimensionality of this dataset, the resulting variably expressed genes were summarized by principle component analysis (PCA), and the first 20 principle components further summarized using Uniform Manifold Approximation and Projection (UMAP) dimensionality reduction. Doublets were assessed using the DoubletFinder (version 2.0.2)^4^ algorithm and few (<10%) doublets were observed outside of the cell population. Clustering was conducted with the FindClusters() function using 20 PCA components and a resolution parameter set to 0.3. Cell cycle analysis was conducted using the CellCycleScoring() in Seurat package with a list of cell cycle markers from Aviv Regev et al’s study^10^.

Differentially expressed genes among clusters and treatments were identified with logFC greater than 0.25 (adjusted p < 0.05) as determined in Wilcoxon rank-sum test from Seurat. The markers for different cell types were retrieved from CellMarker database, which is a manually curated resource of cell markers in human and mouse^11^. The pathway analysis was done using Gene Ontology and canonical pathways in enrichr^12^, with adjusted p < 0.05 as the cutoff for statistically differential pathways.

Cell–cell interactions based on the expression of known ligand–receptor pairs in different cell types were inferred using CellChatDB (v1.0.0)^7^. The total counts of interactions and interaction strengths were calculated using the compareInteractions function in CellChat. The differential edge list was passed through CircosDiff (a wrapper around the R package ‘circlize’) and netVisual_chord_gene in CellChat to filter receptor ligand edges and generate Circos plots.

### Integrated analysis of GBM/HGG and JAK1KO/WT tumors scRNA-seq data

Immune and stromal compartments of the GBM and JAK1KO/WT scRNA-seq data were jointly integrated using rPCA as described above.

### Cytometry by Time-of-flight analysis (CyTOF)

YUMM JAK1/KO and WT tumor cells (0.3×10^6^) were implanted into the flanks of C57BL/6 mice. On day 8 post-inoculation, tumors were harvested from mice at predefined treatment time points.

Tumors were digested using the Tumor Dissociation Kit Mouse (Miltenyi Biotec). Immune cells were isolated using the CD45+ isolation kit (EasySepTM Mouse CD45 Positive Selection Kit, STEMCELL Technologies). A panel of 35 immune markers was used for analysis. Samples were analyzed using the Fluidigm Helios Mass Cytometry System at the UCLA Flow Cytometry core. Manual gating was performed using FlowJo software (version 10.4.2) to identify cells, singlets, and viable CD45+ populations. Data files were analyzed using OMIQ software. Cluster median data were normalized, and a threshold of >0.5 was used to define positive immune markers for cluster identification and annotation.

### Murine RNA Isolation and RNA-seq analysis

For in vitro experiments, melanoma cell lines were seeded of 1.5 × 105 cells per 6-well plate for treatment. After 24 hours, the culture media were replaced with fresh media containing IFNγ (BD Pharmingen, catalog no. 554616) or supernatant of mIL13Rα2 CAR T cells with YUMM2.1 WT tumors. Cells were harvested 8 hours after treatment. The cell pellets were lysed in TRIzol reagent (Invitrogen, catalog no. 15596018) and stored at −80°C until RNA extraction.

For in vivo experiments, engrafted tumors treated with or without mIL13, mIL13-ISG15, or mIL13-mutISG15 CAR T cells were harvested and dissociated into single cells and stored at −80°C prior to RNA extraction.

Total RNA was extracted from mouse tissues using TRIzolTM Reagent (Life Technologies) for transcriptome analyses. RNA integrity, purity, and concentration were evaluated using an Agilent 2100 Bioanalyzer (Agilent Technologies). Nanodrop and Qubit 3.0. RNA-seq libraries were constructed using KAPA mRNA HyperPrep Kit. RNA-seq was performed in the Illumina HiSeq2000 System using 51 bp single-end sequencing. High-quality reads were obtained by trimming the raw reads using fastp (v0.23.3)^13^ and by carrying out FastQC (v0.11.9) for the quality control assessment. The clean reads were aligned to the mouse reference genome GRCm38 with HISAT-STRINGTIE analytic pipeline (HISAT2 v2.1.0; Stringtie v1.3.4) ^14^. The DESeq2 (v.1.26.0)^15^ framework was adopted for differential expression analysis.

### Maintenance of mouse cell lines

The YUMM sublines (YUMM WT and YUMM JAK1/KO) as well as the MC38 cell lines (MC38 WT and MC38 JAK1/KO) were graciously provided by the Antoni Ribas laboratory, located within the comprehensive cancer center at UCLA. Both the YUMM and MC38 sublines were cultured at 37°C with 5% CO2 in DMEM (Invitrogen), supplemented with 10% FBS, 100 U/mL penicillin, 100 μg/mL streptomycin, and 0.25 μg/mL amphotericin B. Regular screenings ensured the absence of Mycoplasma contamination, utilizing the MycoAlert Mycoplasma Detection Kit (Lonza), and authentication tests were periodically conducted. For in vivo experiments, early-passage cell lines (less than 10 passages) were utilized.

### In vivo studies

All experimental procedures involving mice were conducted in accordance with the protocols approved by the City of Hope IACUC. For the subcutaneous models, 3 × 10^6^ YUMM WT and JAK1/KO cells suspended in PBS were injected into the left flanks of 8-10-week-old C57BL/6J mice. After allowing 8 days for tumor establishment, 0.4 × 10^6^ mIL13Rα2 CAR T cells were administered directly into the tumors. Similarly, in the MC38 model, 1 × 10^6^ MC38 WT and JAK1/KO cells were injected into the mice, followed by the same treatment after 8 days. For systemic administration, 5 × 10^6^ mIL13Rα2 CAR T cells were infused via the tail vein on day 8 post-tumor inoculation. Tumor volumes were regularly measured using calipers.

In the orthotopic model, 1 × 10^5^ tumor cells were stereotactically implanted intracranially into the right forebrain of mice. Successful engraftment was confirmed by bioluminescence imaging the day before injecting the CAR T cells. Mice were grouped based on bioluminescence intensity. Fourteen days post-tumor implantation, mice received an intracranial administration of 1 × 10^6^ mIL13Rα2 CAR T cells. Tumor progression was monitored using SPECTRAL LagoX and analyzed with Aura software. Survival outcomes were charted using GraphPad Prism Software (v10).

For combination therapies in flank models, the pretreatment group was administered three doses every other day before CAR T cell therapy and received three additional doses afterward. The simultaneous treatment group was given an initial dose of anti-SPP1(Osteopontin) antibody (bioXcell Clone: 100D3) on the day of CAR T cell therapy, followed by five more doses to ensure parity in antibody delivery across groups, all given intraperitoneally (IP). In the orthotopic model, the combination group received three doses of anti-SPP1 antibody (clone MPIIIB10) prior to the CAR T cell therapy on day 8, continuing with five subsequent doses IP.

All mice were continually observed by the Center for Comparative Medicine at City of Hope for signs of tumor progression and overall survival, with euthanasia applied in adherence to the American Veterinary Medical Association Guidelines.

### Biostatistics

Statistical significance in the in vivo studies was assessed utilizing either a Student t-test for two groups or a one-way ANOVA with Bonferroni correction for three or more groups. Survival analysis was represented via Kaplan-Meier survival curves, with statistical significance determined by the log-rank (Mantel-Cox) test. All statistical analyses were conducted using GraphPad Prism software (v10). Significance levels were denoted as follows: *, P < 0.05; **, P < 0.01; ***, P < 0.001; ****, P < 0.0001.

### YUMM and MC38 sublines expansion

Mouse tumor cells expanded *in vitro* were stained with an unconjugated goat anti-mouse IL13Rα2 (R&D Systems) followed by secondary donkey anti-goat NL637 (R&D Systems). Live murine CAR T cells were stained with CD3 (eBioscience) and CD19 (BD Biosciences) as a surrogate to detect the CAR.

### *ICAM1* engineering of the tumor cells

The plasmid containing the *ICAM1* gene and the packaging plasmids were provided by VectorBuilder. The lentiviral backbone was used for the construction of the expression vector. The cells were then incubated with the lentivirus following the protocol provided by Vectorbuilder. Neomycin (Geneticin) was used for positive selection of the transduced cells. The overexpression of *ICAM1* was confirmed using flow cytometry analysis.

### In vitro verification of YUMM and MC38 sublines

To fully characterize the differential IFN-induced gene expression, the engineered YUMM JAK1/KO and YUMM-WT as well as MC38 JAK1/KO and MC38 WT were exposed to supernatants derived from CAR T cell mediated cytotoxicity with matched YUMM and MC38 WT tumors. The cells, subsequently, were sent to bulk RNA-sequencing for transcriptional analysis.

### Murine CAR T cells

The murine IL13Rα2 CAR was engineered within an MSCV retroviral backbone (Addgene), comprising the murine IL13 extracellular domain, murine CD8 hinge, murine CD8 transmembrane domain, and intracellular murine 4-1BB costimulatory and murine CD3ζ signaling domains. A truncated murine CD19 was inserted downstream via a T2A ribosomal skip as a transduction marker. The resulting plasmid was transfected into PlatE cells (courtesy of Dr. Zuoming Sun’s lab) using Fugene (Promega). After 48 hours, the supernatant was harvested and filtered through a 0.2-μm filter, then aliquoted and stored frozen until transduction.

Murine T cells were isolated from spleens of naïve C57BL/6J mice or using the EasySep Mouse T cell Isolation Kit (STEMCELL Technologies) and activated with Dynabead Mouse T-Activator CD3/CD28 beads (Gibco) at a 1:1 ratio. Transduction of T cells occurred on RetroNectin-coated plates (Takara Bio) using retrovirus-containing supernatants (as described above). Cells were then expanded for 4 days in RPMI 1640 (Lonza) supplemented with 10% FBS (Hyclone Laboratories), 55 mmol/L 2-mercaptoethanol (Gibco), 50 U/mL recombinant human IL2 (Novartis), and 10 ng/mL recombinant murine IL7 (PeproTech). Prior to in vitro and in vivo experiments, beads were magnetically separated from T cells and CAR expression was assessed via flow cytometry.

### In vitro cytotoxicity using Xcelligence assay

The assay was performed following a standardized protocol. On Day 1, the plate map was set up in the Xcelligence software based on the experimental design. Target cells were dissociated using trypsin or scraped to remove enzymes, resuspended in assay media, and passed through a 40 μm cell strainer to eliminate clumps. Cell counts were determined using the Muse Cell Analyzer, and cells were resuspended at the desired concentration. A total of 50 μL of assay media was added to each well of the E-plate, and background signal was recorded on the Xcelligence instrument, ensuring no bubbles were present. The plate was then removed, and 100 μL of the target cell suspension was added to each well. Cells were allowed to settle for 45 minutes at room temperature before placing the plate back on the instrument to minimize edge effects and ensure accurate readings. Target cells were cultured for 24 hours at 37°C in a humidified incubator with 5% CO₂. On Day 2, effector cells were prepared at a concentration of 1.0 × 10⁶ cells/mL for each cell line, followed by serial dilutions to achieve the desired effector-to-target (E:T) ratio. The Xcelligence experiment was paused, and 100 μL of the prepared effector cell suspension was added to each well in a tissue culture hood. Cells were allowed to settle for 45 minutes at room temperature before resuming the experiment on the Xcelligence instrument.

### Immunofluorescence staining

Immunofluorescence was done on 5 µm-thick sections of FFPE (formalin-fixed paraffin-embedded) specimens placed on positively charged glass slides. The slides were deparaffinized in xylene, rehydrated in an ethanol gradient, and underwent antigen retrieval (10 min, 110-116 o C) in citrate-based antigen unmasking solution (#H-3300, Vector Laboratories) using a pressure cooker. The slides were then washed and incubated with carbohydrate-free blocking solution (#SP-5040, Vector Laboratories) prior to staining. The slides were then incubated with anti-COL1A1 antibody (E3E1X, Cell Signaling, #66948), anti-SPP1 (#AF1433, R&D) and anti-CD68 (EPR20545, Abcam, #ab213363) overnight at 4 °C, followed by incubation with donkey anti-mouse Alexa Fluor 555 (#A-31570), donkey anti-goat Alexa Fluor 488 (#A-11055), donkey anti-rabbit Alexa Fluor 647 (#A-31573) and Hoechst 33342 (#H3570, Thermofisher Scientific) for 1 hour (RT). The slides were then coverslipped and scanned using a Zeiss LSM880 confocal microscope. Murine tissues were incubated with anti-COL1A1 antibody (E8F4L, Cell Signaling, #72026), anti-SPP1 (#AF808, R&D) and anti-CD68 (FA-11, Invitrogen, #14-0681-82), followed by incubation with donkey anti-rat Alexa Fluor 594 (#A-21209), donkey anti-mouse Alexa Fluor 647 (#A-31571), donkey ant-rabbit Alexa Fluor 647 (#A-31573) and Hoechst 33342 (#H3570).

### Data and code availability

Raw and processed GBM/HGG scRNA-seq data are deposited at the Gene Expression Omnibus (GEO) under accession number GSE290291 and mouse data under accession GSE289327. Custom scripts used for data analysis are available on GitHub at https://github.com/Banovich-Lab/HGG_SPP1.

---

**Extended Fig 1.**
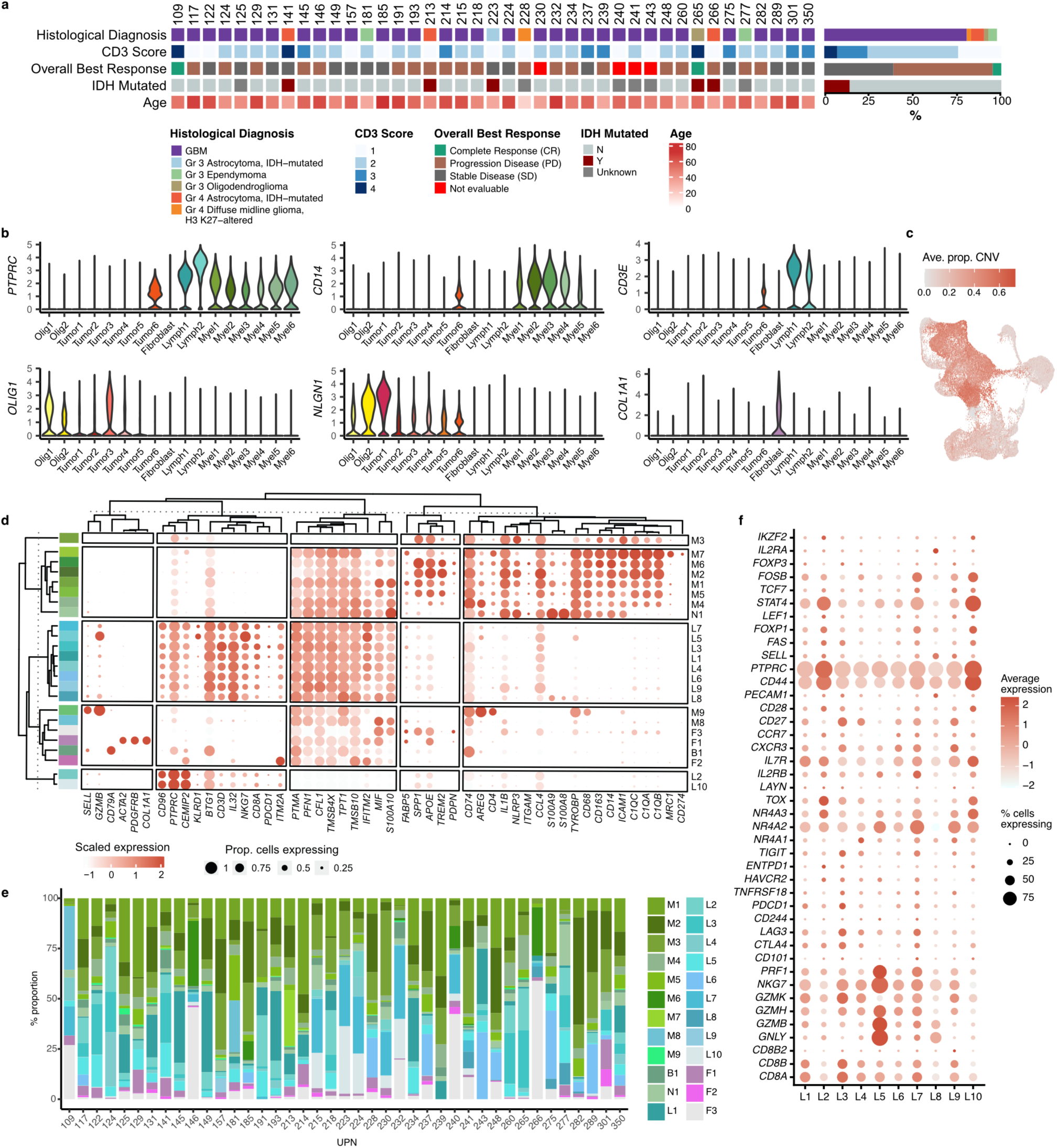
Demographic and clinical characterization of tumor samples included from 41 patients with GBM/HGG 1. a, Demographic and clinical characteristics of tumors from 41 patients with high-grade glioma. **b,** Expression of global marker features across GBM/HGG tumor clusters. **c,** Average proportion of copy number alterations for each cell in scRNA-seq data of GBM/HGG tumors. **d,** Expression of canonical myeloid, lymphoid, and fibroblast marker features across cell types in the TME of GBM/HGG tumors. **e,** Proportions of cells annotated for each cluster in the TME of GBM/HGG tumor samples from the 41 donors. **f,** Expression of canonical T cell markers in the lymphoid subclusters of GBM/HGG tumors.

**Extended Fig. 2.**
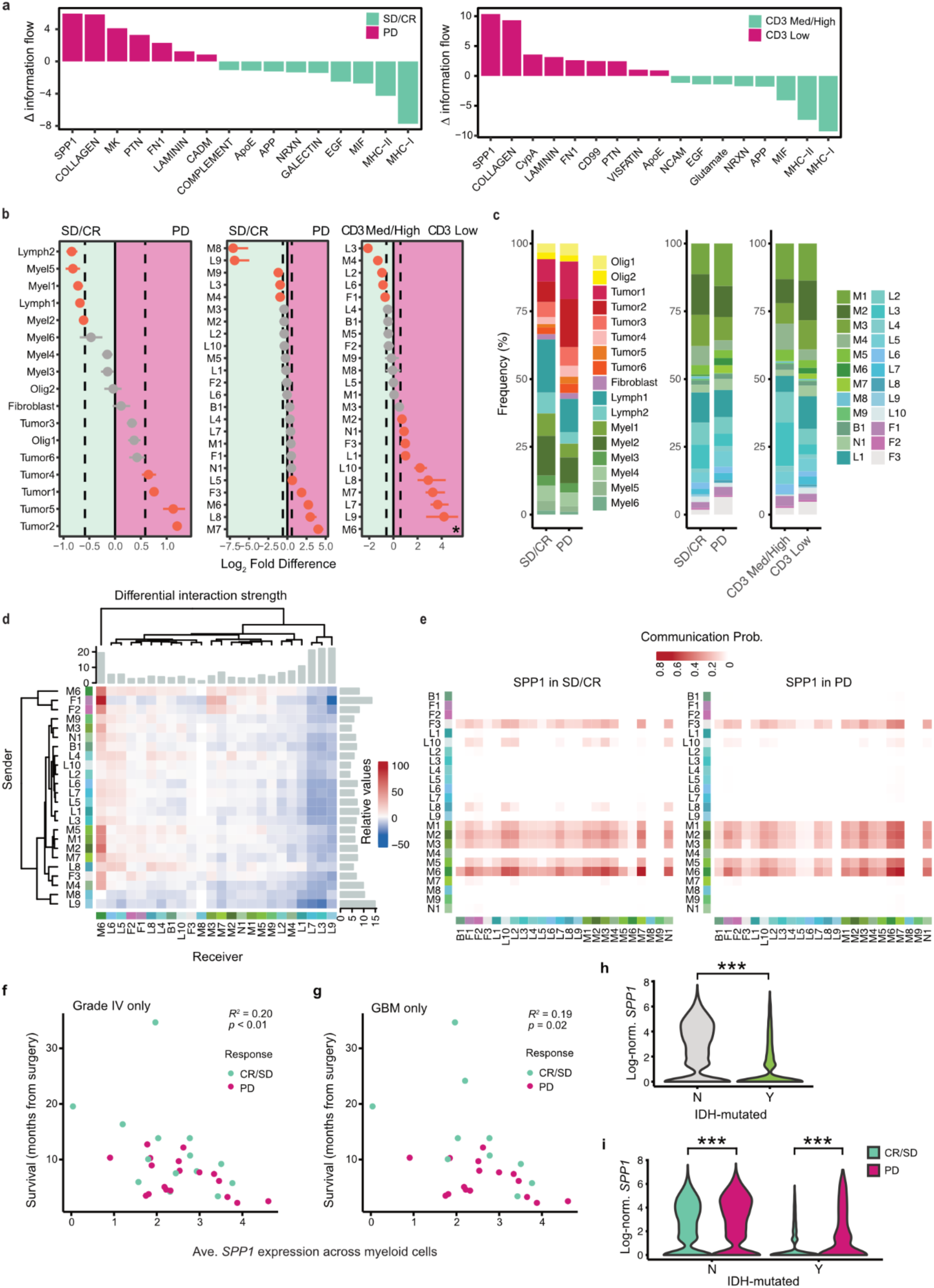
Responder-versus-non-responder TME signaling differences and SPP1 correlations in GBM/HGG. a, Top signaling pathways differentially regulated between SD/CR and PD GBM/HGG tumors (left) and CD3-med/high and CD3-low tumors (right). **b,** Differential abundance of all cell types between SD/CR and PD GBM/HGG tumors (left), the TME of SD/CR and PD GBM/HGG tumors (middle), and the TME of CD3-med/high and CD3-low GBM/HGG tumors (right). Log_2_ fold-differences with 95% confidence intervals are presented. Results for cell types with significant differences based on a permutation p-value are highlighted in red. In the CD3-med/high vs. CD3-low comparison, M6 was only present in CD3-low (indicated with an asterisk). **c,** Proportions of all cell types in SD/CR GBM/HGG tumors (left) as well as the TME of SD/CR and PD (middle) and CD3-med/high and CD3-low GBM/HGG tumors (right). **d,** Differential signaling strength between all cell types in the TME of SD/CR and PD GBM/HGG tumors. Positive values indicate increased signaling in PD. Barplots depict the total signaling strength for each sender and receiver. **e,** *SPP1* signaling communication probability between all pairs of cell types in the TME of SD/CR (left) and PD (right) GBM/HGG tumors. **f,** Relationships between survival and average *SPP1* expression in the myeloid fraction of Grade IV GBM/HGG tumors (*R*^2^ and p-value are indicated). **g,** Relationships between survival and average *SPP1* expression in the myeloid fraction of GBM tumors (*R*^2^ and p-value are indicated). **h,** *SPP1* expression in the myeloid fraction of IDH wildtype and IDH-mutated tumors. **i,** *SPP1* expression in the myeloid fraction of IDH wildtype and IDH-mutated GBM/HGG tumors by response to CAR T therapy. In (g) and (h), *** indicates adjusted p-value < 0.01.

**Extended Fig. 3.**
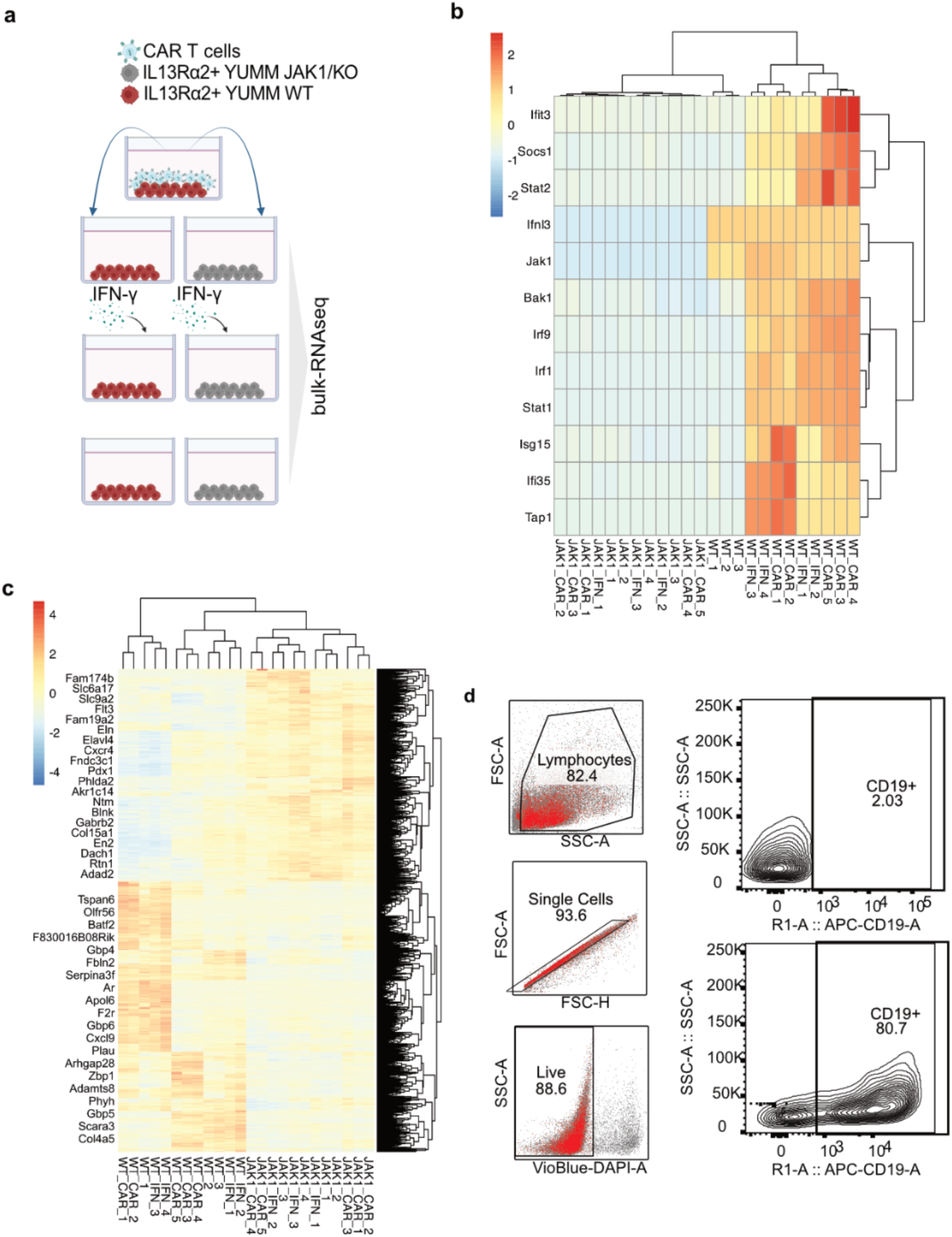
Transcriptional alterations of YUMM WT and YUMM JAK1/KO tumor cells post exposure to mIL13CAR T cell-tumor cell interaction. a, Schematic illustration demonstrating the verification of YUMM WT and YUMM JAK1/KO tumors. mIL13CAR T cells were exposed to WT tumor cells, and the supernatant from this exposure was added to YUMM WT and JAK1/KO tumors. Tumors post-exposure to media from CAR T cell-tumor cell interactions were collected for bulk deconvolution analysis. **b, c,** The analysis revealed altered mIL13Rα2 CAR T cell-induced gene transcription in JAK1/KO YUMM melanoma cells compared to WT, specifically in the IFN signaling pathway and broader holistic pathways. Heatmap displaying the change in gene expression due to mIL13Rα2 CAR T cell treatment. Genes were sorted by the most highly enriched pathways. **d,** Background gating for separating out engineered mIL13Rα2 CAR T cells.

**Extended Fig. 4.**
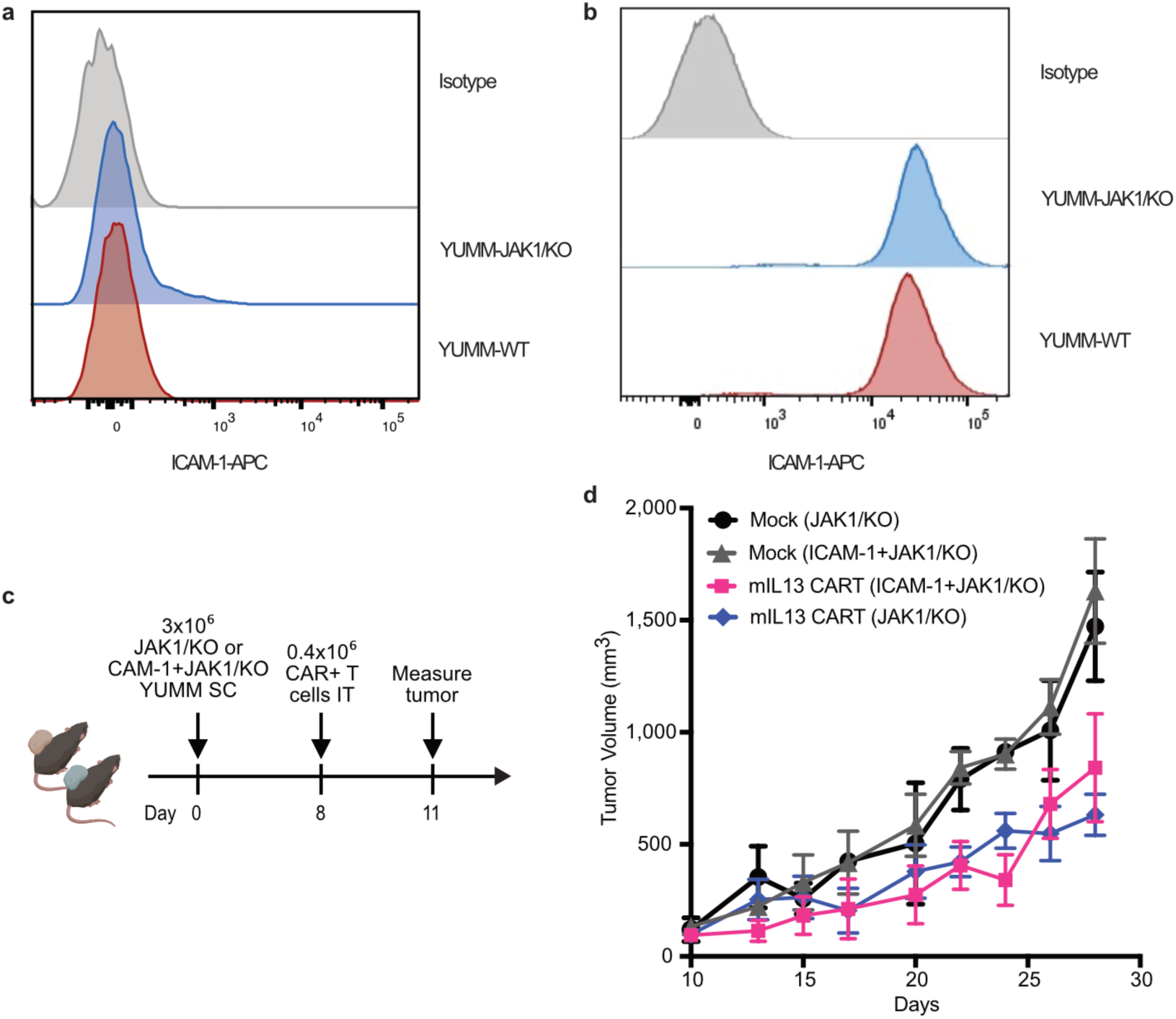
*ICAM1* expression and tumor growth dynamics in JAK1/KO melanoma models treated with mIL13R CAR T cells. a-b, Flow cytometry analysis of *ICAM1* surface expression in YUMM-JAK1/KO and YUMM-WT melanoma cells before and after *ICAM1* transduction. Histogram showing *ICAM1* expression in YUMM-JAK1/KO (blue) compared to YUMM-WT (red), with isotype control (gray). **c,** Schematic representation of the experimental setup. JAK1/KO or *ICAM1*+JAK1/KO YUMM melanoma cells (3×10⁶) were subcutaneously implanted in mice (Day 0) (n=5 per arm). On Day 8, mice received 0.4×10⁶ mIL13Rα2 CAR T cells IT and tumor measurement started on Day 11. **c,** Tumor growth curves in mice implanted with JAK1/KO or *ICAM1*+JAK1/KO tumors and treated with either mock or mIL13Rα2 CAR T cells. No significant changes were detected in tumors growth comparing JAK1/KO to *ICAM1*+JAK1/KO tumors treated with mIL13Rα2 CAR T cells.

**Extended Fig. 5.**
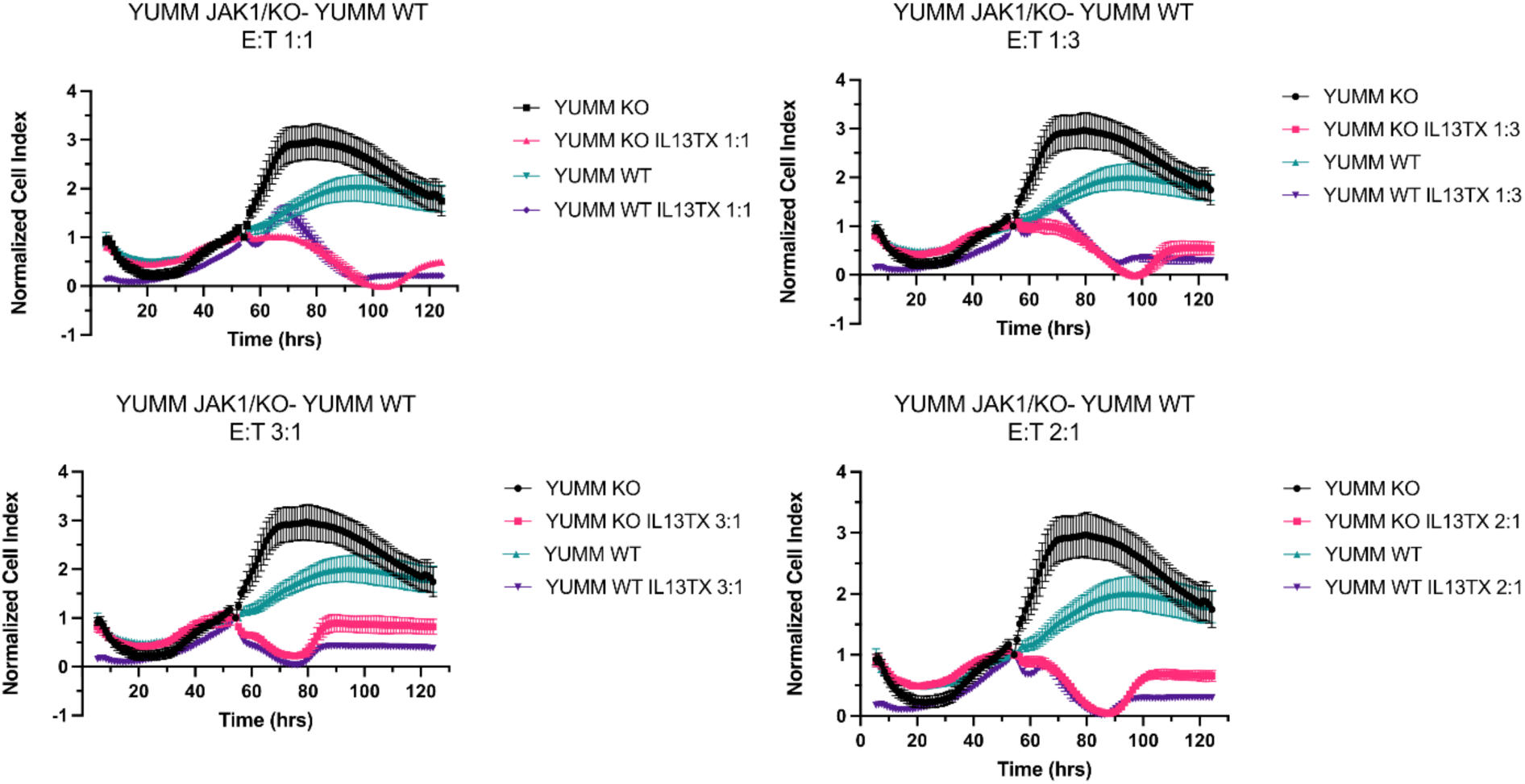
Comparable mIL13 CAR T cell cytotoxicity in YUMM JAK1/KO and WT melanoma cells across multiple effector-to-target (E:T) ratios. Real-time impedance-based cytotoxicity assay measuring IL13Rα2-targeted CAR T cell-mediated killing of YUMM JAK1/KO and YUMM WT melanoma cells at different E:T ratios (1:1, 1:3, 2:1, and 3:1). The normalized cell index represents tumor cell viability over time. IL13 CAR T cells effectively reduced tumor cell viability in both JAK1/KO and WT YUMM cells indicating comparable in vitro cytotoxicity.

**Extended Fig. 6.**
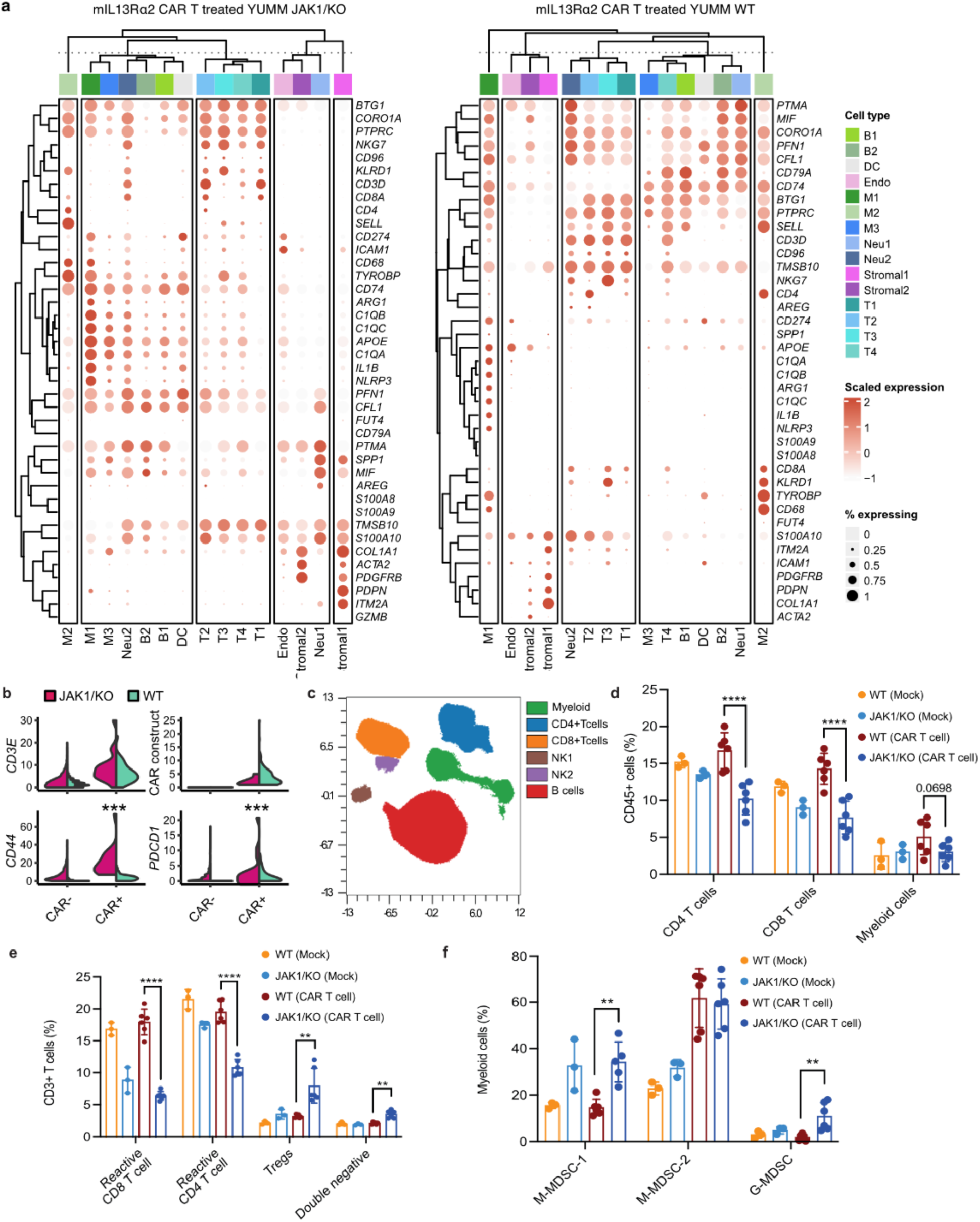
Immune landscape of JAK1/KO YUMM and WT YUMM tumors after CAR T cell therapy. a, Expression of canonical myeloid, lymphoid, and fibroblast markers across cell types in the TME of mIL13Rα2 CAR T treated YUMM JAK1/KO and WT tumors. **b**, Expression of *CD44* and *PDCD1* (PD-1) in CAR positive and CAR negative T cells in JAK1/KO and WT YUMM tumors. *** denotes differential expression (adj. p < 0.001). **c**, t-SNE plot of CyTOF data showing CD45+ immune cell clustering in tumors, with major subsets including myeloid cells, CD4+ T cells, CD8+ T cells, NK1, NK2, and B cells(n=5 in treated arms, n=3 in control arms) **d,** Quantification of CD45+ immune subsets as a percentage of total immune cells in tumors under different conditions: WT (Mock), JAK1KO (Mock), WT (CAR T cell-treated), and JAK1/KO (CAR T cell-treated). JAK1/KO YUMM tumors exhibit significantly reduced CD4+ and CD8+ T cell infiltration after CAR T cell therapy compared to WT tumors (****p < 0.0001). **e,** CD3+ T cell subpopulations show a marked decrease in reactive CD8+ and CD4+ T cells in YUMM JAK1/KO tumors, alongside significant increase of regulatory T cells and double-negative T cells compared to YUMM WT (** p < 0.01, *** p < 0.001, **** p < 0.0001). **f,** Myeloid-derived suppressor cell (MDSC) composition in tumors, demonstrating an increased presence of monocytic-MDSC (M-MDSC1, M-MDSC2) and granulocytic-MDSC (G-MDSC) in YUMM JAK1/KO tumors compared to YUMM WT tumors (**p < 0.01).

**Extended Fig. 7.**
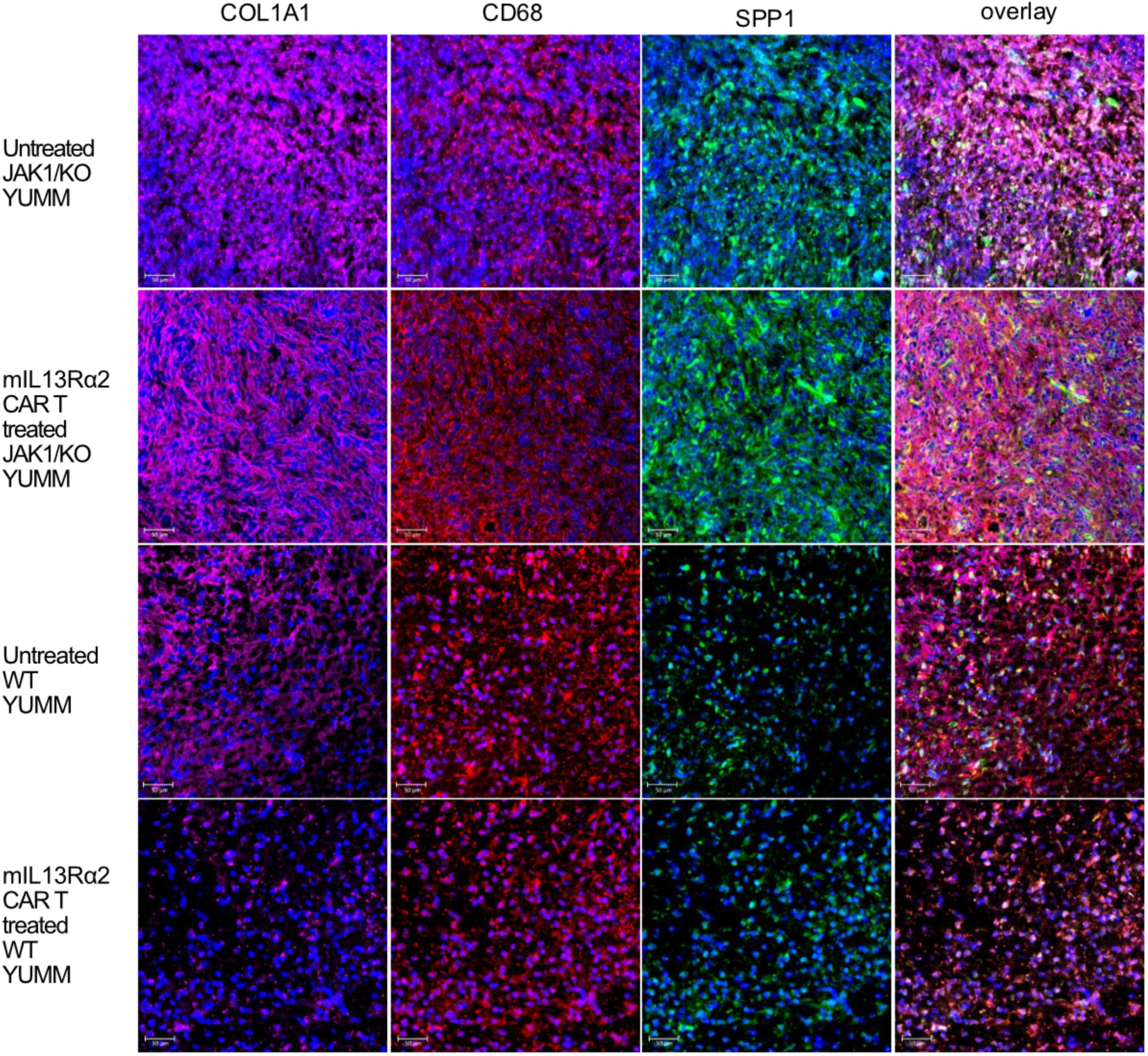
**Immunofluorescence analysis in YUMM JAK1/KO versus WT YUMM tumors after mIL13Rα2 CAR T-cell therapy**. Representative confocal micrographs of resected subcutaneous tumors harvested on day 10 post-treatment. Sections were stained for COL1A1 (purple), CD68 (red), SPP1 (green) and nuclei (DAPI, blue), Scale bars, 100 μm.

**Extended Fig. 8.**
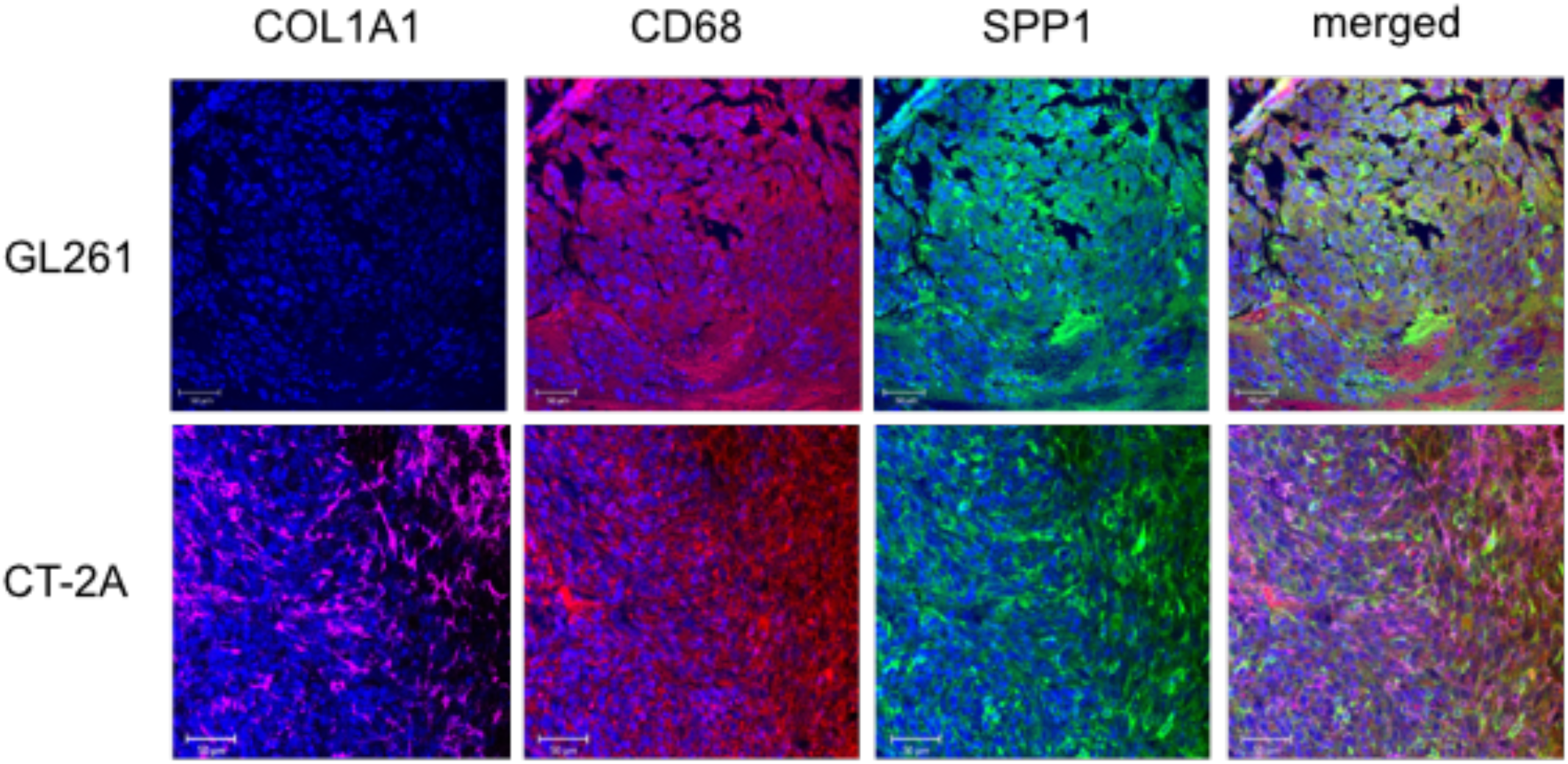
**Immunofluorcent staining for COL1A1, CD68, and SPP1 in GL261 and CT-2A mouse glioma models**. Representative brain sections from orthotopic GL261 and CT-2A gliomas stained for COL1A1 (purple), CD68 (red), SPP1 (green) and nuclei (DAPI, blue); Scale bars 50 μm.

## Notes

### Competing Interest Statement

The authors have declared no competing interest.

### Summary of Updates

There have been some errors and inconsistencies in figure labels and text. We are uploading a revision with corrections to the mentioned areas.

